# Phosphorylation bar-coding of Free Fatty Acid receptor 2 is generated in a tissue-specific manner

**DOI:** 10.1101/2023.09.01.555873

**Authors:** Natasja Barki, Laura Jenkins, Sara Marsango, Domonkos Dedeo, Daniele Bolognini, Louis Dwomoh, Aisha M. Abdelmalik, Margaret Nilsen, Manon Stoffels, Falko Nagel, Stefan Schulz, Andrew B. Tobin, Graeme Milligan

## Abstract

Free Fatty Acid receptor 2 (FFA2) is activated by short-chain fatty acids and expressed widely, including in white adipocytes and various immune and enteroendocrine cells. Using both wild type human FFA2 and a Designer Receptor Exclusively Activated by Designer Drugs (DREADD) variant we explored the activation and phosphorylation profile of the receptor, both in heterologous cell lines and in tissues from transgenic knock-in mouse lines expressing either human FFA2 or the FFA2-DREADD. FFA2 phospho-site specific antisera targeting either pSer^296^/pSer^297^ or pThr^306^/pThr^310^ provided sensitive biomarkers of both constitutive and agonist-mediated phosphorylation as well as an effective means to visualise agonist-activated receptors *in situ*. In white adipose tissue phosphorylation of residues Ser^296^/Ser^297^ was enhanced upon agonist activation whilst Thr^306^/Thr^310^ did not become phosphorylated. By contrast, in immune cells from Peyer’s patches Thr^306^/Thr^310^ become phosphorylated in a strictly agonist-dependent fashion whilst in enteroendocrine cells of the colon both Ser^296^/Ser^297^ and Thr^306^/Thr^310^ were poorly phosphorylated. The concept of phosphorylation bar-coding has centred to date on the potential for different agonists to promote distinct receptor phosphorylation patterns. Here we demonstrate that this occurs for the same agonist-receptor pairing in different patho-physiologically relevant target tissues. This may underpin why a single G protein-coupled receptor can generate different functional outcomes in a tissue-specific manner.

**Significance Statement:** The concept that agonist-occupancy of a G protein-coupled receptor can result in distinct patterns of phosphorylation of residues on the intracellular elements of the receptor in different tissues is referred to ‘bar-coding’. This has been challenging to demonstrate conclusively in native tissues. We now show this to be the case by using tissues from transgenic knock-in mouse lines expressing either wild type or a DREADD variant of human Free Fatty Acid Receptor 2 and a pair of phospho-site specific antisera. Clear differences in the pattern of phosphorylation of the receptor induced by the same ligand were observed in white adipose tissue and immune cells derived from Peyer’s patches. These outcomes provide direct evidence in tissues, at endogenous expression levels, of a well promoted hypothesis.

## Introduction

G protein-coupled receptors (GPCRs) routinely are constitutively phosphorylated or become phosphorylated on serine and threonine residues, located either within the 3^rd^ intracellular loop or the C-terminal tail, following exposure to activating agonist ligands (1). This is generally accompanied by new or enhanced interactions with arrestin adapter proteins. Canonically this results in receptor desensitization because the positioning of the arrestin precludes simultaneous interactions of the GPCR with a heterotrimeric guanine nucleotide binding (G)-protein and hence ‘arrests’ further G protein activation (2). The earliest studies on GPCR phosphorylation determined that receptors exist in multiply phosphorylated states where numerous kinases, that include members of the G-protein receptor kinase (GRK) family, second messenger-regulated kinases and even those of the casein kinase family are involved (3-5). Using mass spectrometry approaches and phosphosite-specific antibodies it emerged that the pattern of receptor phosphorylation was distinct among cell types and tissues (6-8). This led to the notion that the tissue-specific signalling output of GPCRs might be determined, at least in part, by the pattern of receptor phosphorylation – a hypothesis coined the phosphorylation barcode (7, 9). This notion has recently been taken a step further with appreciation that the conformation adopted by an arrestin on interaction with the phosphorylated form of a GPCR might be affected by the pattern of receptor phosphorylation and in turn this may affect the signalling outputs mediated by arrestins (10). The challenge in fully appreciating the impact of the receptor phosphorylation barcode on the physiological activity of GPCRs is in the identification of the receptor-phosphorylation patterns in native tissues. Here we address this issue by use of antibodies raised against specific phosphorylation sites within the free fatty acid receptor 2 (FFA2).

A number of GPCRs are able to bind and respond to short chain fatty acids (SCFAs) that are generated in prodigious quantities by fermentation of dietary fibre by the gut microbiota (11,12). The most studied and best characterized of these is FFA2 (also designated GPR43) (13-15). This receptor is expressed widely, including by a range of immune cells, adipocytes, enteroendocrine and pancreatic cells (13-15). This distribution has led to studies centred on its potential role at the interface of immune cell function and metabolism (11,12,16), as well as other potential roles in regulation of gut mucosal barrier permeability (11,12) and the suppression of bacterial and viral infections (17, 18). In addition to being widely expressed FFA2 is, at least when expressed in simple heterologous cell lines, able to couple to of a range of different heterotrimeric G proteins from each of the G_i_, G_q_ and G_12_/G_13_ families (19, 20). Despite this, however, studies suggest more selective activation of signalling pathways are induced by stimulation of FFA2 in different native cells and tissues. For example, in white adipocytes stimulation of FFA2 is clearly anti-lipolytic, an effect mediated via pertussis toxin-sensitive G_i_-proteins and reduction in cellular cAMP levels (21), whilst in GLP-1 positive enteroendocrine cells activation of FFA2 promotes release of this incretin hormone in a Ca^2+^ and G_q_-mediated manner (21).

A challenge in studying the molecular basis of effects of SCFAs in native cells and tissues is that the pharmacology of the various GPCRs that respond to these ligands is very limited (22,23) and there is a particular dearth of antagonist ligands for anything other than the human ortholog of FFA2 (22,23). DREADD forms of GPCRs contain mutations that eliminate binding and responsiveness to endogenously generated orthosteric agonists whilst, in parallel, allowing binding and activation by defined, but non-endogenously produced, ligands that do not activate the wild-type form of the receptor (24-26). Such DREADDs have become important tools to probe receptor function in the context of cell and tissue physiology. Here, in addition to studying wild type human FFA2 we also employed a DREADD form of human FFA2 (hFFA2-DREADD) that we have previously generated (27) and characterized extensively (20, 28-29). This has the benefit of providing both small, highly selective, water-soluble orthosteric agonists for FFA2 and, when using the human ortholog of FFA2, there are high affinity and well characterized antagonists (22,23, 30) that can be used to further define on-target specificity of effects that are observed.

## Results

### Generation and characterization of phospho-site specific antisera to identify activated hFFA2-DREADD

We performed mass spectrometry on a form of hFFA2-DREADD with C-terminally linked enhanced Yellow Fluorescent Protein (hFFA2-DREADD-eYFP) following its doxycycline-induced expression in Flp-In T-REx 293 cells. These studies identified phosphorylation at residue Ser^297^ in both the basal state and after addition of the orthosteric hFFA2-DREADD agonists (2E,4E)-hexa-2,4-dienoic acid (sorbic acid) (20, 27) or 4- methoxy-3-methyl-benzoic acid (MOMBA) (28). In addition, the adjacent residue Ser^296^ was also observed to be phosphorylated only after agonist treatment (**Figure 1**). Based on these outcomes we generated an antiserum anticipated to identify hFFA2-DREADD when either pSer^296^, pSer^297^, or both these amino acids, were phosphorylated (**Figures 2A, 2B**). Ser^296^ and Ser^297^ are within the intracellular C-terminal tail of the receptor (**Figure 2A**) and within this region there are other potential phospho-acceptor sites. These include Thr^306^ and Thr^310^ as well as Ser^324^ and Ser^325^ (**Figure 2A**). Although we did not obtain clear evidence of either basal or DREADD agonist-induced phosphorylation of these residues from the mass spectrometry studies we also generated antisera potentially able to identify either pThr^306^/pThr^310^ hFFA2-DREADD (**Figures 2A, 2B**) or pSer^324^/pSer^325^ hFFA2-DREADD (**not shown**).

**Figure 1.**
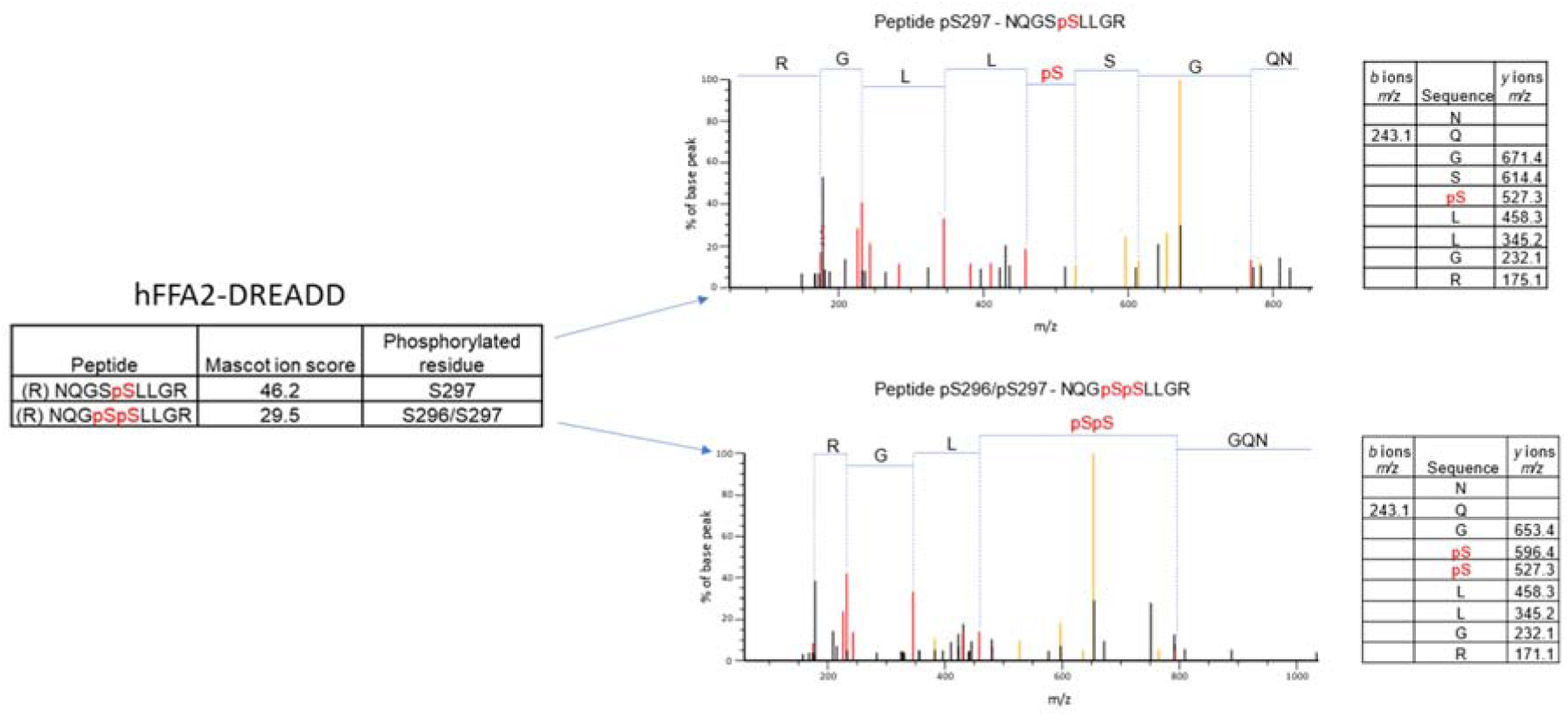
Mass spectrometry analysis of hFFA2-DREADD-eYFP identifies basal phosphorylation of Ser^297^ and agonist-promoted phosphorylation of Ser^296^. Mass spectrometry analysis was conducted on samples isolated from Flp-In T-REx 293 cells in which expression of hFFA2-DREADD-eYFP had been induced. Experiments were performed on vehicle and MOMBA-treated (100 μM, 5 min) cells as detailed in Experimental. LC-MS/MS identified Ser^297^ as being phosphorylated constitutively, and Ser^296/297^ as being phosphorylated by sorbic acid or MOMBA. Composite outcomes of a series of independent experiments are combined. Fragmentation tables associated with phosphorylated peptides are shown. Phosphorylated residues are highlighted in *red*.

**Figure 2.**
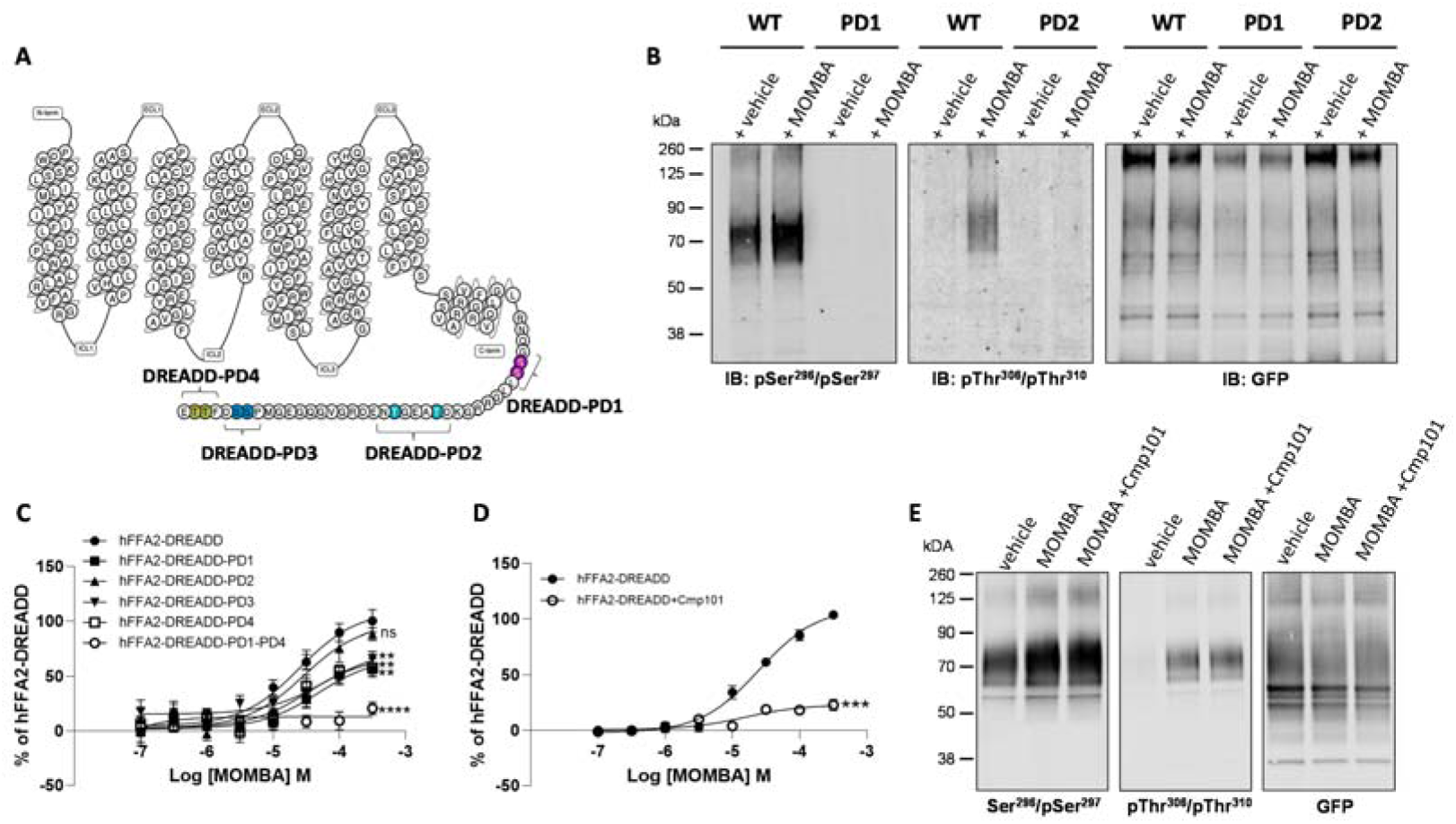
Characteristics of putative pSer^296^ /pSer^297^ and pThr^306^ /pThr^310^ hFFA2- antisera and the effect of potential phospho-acceptor site mutations on agonistinduced arrestin-3 interactions. The primary amino acid sequence of hFFA2 is shown (**A**). Residues altered to generate the DREADD variant are in red (Cys^141^ Gly, His^242^ Gln). Phospho-deficient (PD) hFFA2- DREADD variants were generated by replacing serine 296 and serine 297 (purple, hFFA2- DREADD-PD1), threonine 306 and threonine 310 (light blue, hFFA2-DREADD-PD2), serine 324 and serine 325 (dark blue, hFFA2-DREADD-PD3) or threonine 328 and 329 (yellow, hFFA2-DREADD-PD4) with alanine. In addition, hFFA2-DREADD-PD1-4 was generated by combining all these alterations, **B.** The ability of putative pSer^296^ /pSer^297^ and pThr^306^ /Thr^310^ antisera to identify wild type and either PD1 or PD2 forms of hFFA2- DREADD with and without treatment of cells expressing the various forms with MOMBA is shown and, as a control, anti-GFP immunoblotting of equivalent samples is illustrated. **C**. The ability of varying concentrations of MOMBA to promote interaction of arrestin-3 with hFFA2-DREADD and each of the DREADD-PD mutants, is illustrated. Each of the DREADD-PD variants, except hFFA2-DREADD-PD2 (ns), were less effective in promoting interactions in response to MOMBA (** p < 0.01, **** p < 0.0001). **D**. The effect of the GRK2/3 inhibitor compound 101 on the capacity of MOMBA to promote recruitment of arrestin-3 to wild type hFFA2-DREADD is shown (*** p < 0.001). Significance in **C** and **D** were assessed by one-way ANOVA followed by Dunnett’s multiple comparisons test. **E**. The effect of compound 101 on detection of hFFA2- DREADD-eYFP by each of pSer^296^ /pSer^297^, pThr^306^ /Thr^310^ and anti-GFP antisera is shown. Data are representative (**B, E**) or show means +/- SEM (**C, D**) of at least 3 independent experiments.

To initially assess these antisera we performed immunoblots using membrane preparations from the same Flp-In T-REx 293 cells that had been induced to express hFFA2-DREADD-eYFP. The potential pSer^296^/pSer^297^ antiserum was able to identify a diffuse set of polypeptides centred at 70 kDa in preparations generated from vehicle-treated cells (**Figure 2B**) and the intensity of staining was increased in samples derived from cells after exposure to the hFFA2-DREADD agonist MOMBA (**Figure 2B**). By contrast when samples were produced from a Flp-In T-REx 293 cell line induced to express a variant of hFFA2- DREADD-eYFP in which both Ser^296^ and Ser^297^ were converted to Ala (hFFA2-DREADD- PD1) **(Figures 2A, 2B**) the potential pSer^296^/pSer^297^ antiserum was unable to identify the receptor protein either with or without exposure of the cells to MOMBA (**Figure 2B**). Similar studies were performed with the putative pThr^306^/pThr^310^ antiserum on samples induced to express hFFA2-DREADD-eYFP. In this case there was no significant detection of the receptor construct in samples from vehicle-treated cells (**Figure 2B**), however, after exposure to MOMBA there was also strong identification of diffuse polypeptide(s) centred at 70 kDa corresponding to hFFA2-DREADD-eYFP (**Figure 2B**). Such staining was absent, however, when a variant hFFA2-DREADD-eYFP, in which in this case both Thr^306^ and Thr^310^ were altered to Ala (hFFA2-DREADD-PD2), was induced (**Figures 2A, 2B**). To ensure that the lack of recognition by the potential phospho-site specific antisera was not simply due to poor expression of either the hFFA2-DREADD-PD1 or hFFA2-DREADD-PD2 variants we also immunoblotted equivalent samples with an anti-GFP antiserum (that also identifies eYFP).

This showed similar levels of detection of the approximately 70kDa polypeptide(s) in all samples (**Figure 2B**). Although as noted earlier we did attempt to generate a potential pSer^324^/pSer^325^ antiserum, we were unable to detect the receptor using this in samples containing hFFA2-DREADD-eYFP exposed to either MOMBA or vehicle (**data not shown**). As such, this was not explored further. However, as we also generated both a hFFA2- DREADD-PD3 variant, in which both Ser^324^ and Ser^325^ were altered to alanine (**Figure 2A**), and a hFFA2-DREADD-PD4 variant (**Figure 2A**) in which both Thr^328^ and Thr^329^ were altered to alanine, we assessed the effects of these changes on the capacity of MOMBA to promote interactions of wild type and the phospho-deficient (PD)-variants of hFFA2- DREADD-eYFP with arrestin-3. MOMBA promoted such interactions with the intact DREADD receptor construct in a concentration-dependent manner with pEC_50_ = 4.62 +/- 0.13 M (mean +/- SEM, n = 3). The maximal effect, but not measured potency, of MOMBA was reduced for all the PD variants except hFFA2-DREADD-PD2 (**Figure 2C**), but in no case was the effect on maximal signal reduced by more than 50%. We hence combined these PD variants to produce a form of the receptor in which all potential phospho-acceptor sites in the C-terminal tail were altered to alanines (hFFA2-DREADD-PD1-4). This variant was poorly able to recruit arrestin-3 (20.8 +/- 2.3% of wild type receptor, mean +/- S.D, n = 3) in response to addition of MOMBA (**Figure 2C**). It is likely that GRK2 and/or GRK3 played a key role in allowing MOMBA-induced interaction with arrestin-3 because this was greatly reduced when cells expressing the wild-type form of the receptor construct were pre-treated with the GRK2/GRK3 selective inhibitor compound 101 (31) (**Figure 2D**). To extend this analysis we pre-treated cells induced to express hFFA2-DREADD-eYFP with either vehicle or compound 101 and then, after addition of MOMBA, assessed receptor phosphorylation as detected by the putative pThr^306^/pThr^310^ and pSer^296^/pSer^297^ antisera. Pretreatment with compound 101 did not substantially reduce MOMBA-mediated identification by either the pThr^306^/pThr^310^ antiserum or the pSer^296^/pSer^297^ antiserum (**Figure 2E**). This suggests that phosphorylation of these sites are not controlled by GRK2/3 and that these are not, at least in isolation, sufficient to define interactions of the receptor with arrestin-3.

### Effects of MOMBA on antisera recognition are both on-target and reflect receptor phosphorylation

The selective effect of the hFFA2-DREADD agonist MOMBA in promoting enhanced phosphorylation of Ser^296^/Ser^297^ and in allowing phosphorylation of Thr^306^/Thr^310^ was further substantiated because both these effects of MOMBA were re-capitulated by the second hFFA2-DREADD specific agonist, sorbic acid (20, 27) (**Figure 3A**). By contrast, propionic acid (C3) which, as a SCFA, is an endogenous activator of wild-type hFFA2 but does not activate hFFA2-DREADD (20), was unable to modulate the basal levels of phosphorylation detected by the pSer^296^/pSer^297^ antiserum or to promote detection of the receptor by the pThr^306^/pThr^310^ antiserum (**Figure 3A**). Use of the anti-GFP antiserum confirmed similar levels of receptor in the vehicle and C3-treated samples as in those treated with MOMBA or sorbic acid and, in addition, the lack of expression of the receptor construct prior to doxycycline-induction (**Figure 3A**).

**Figure 3.**
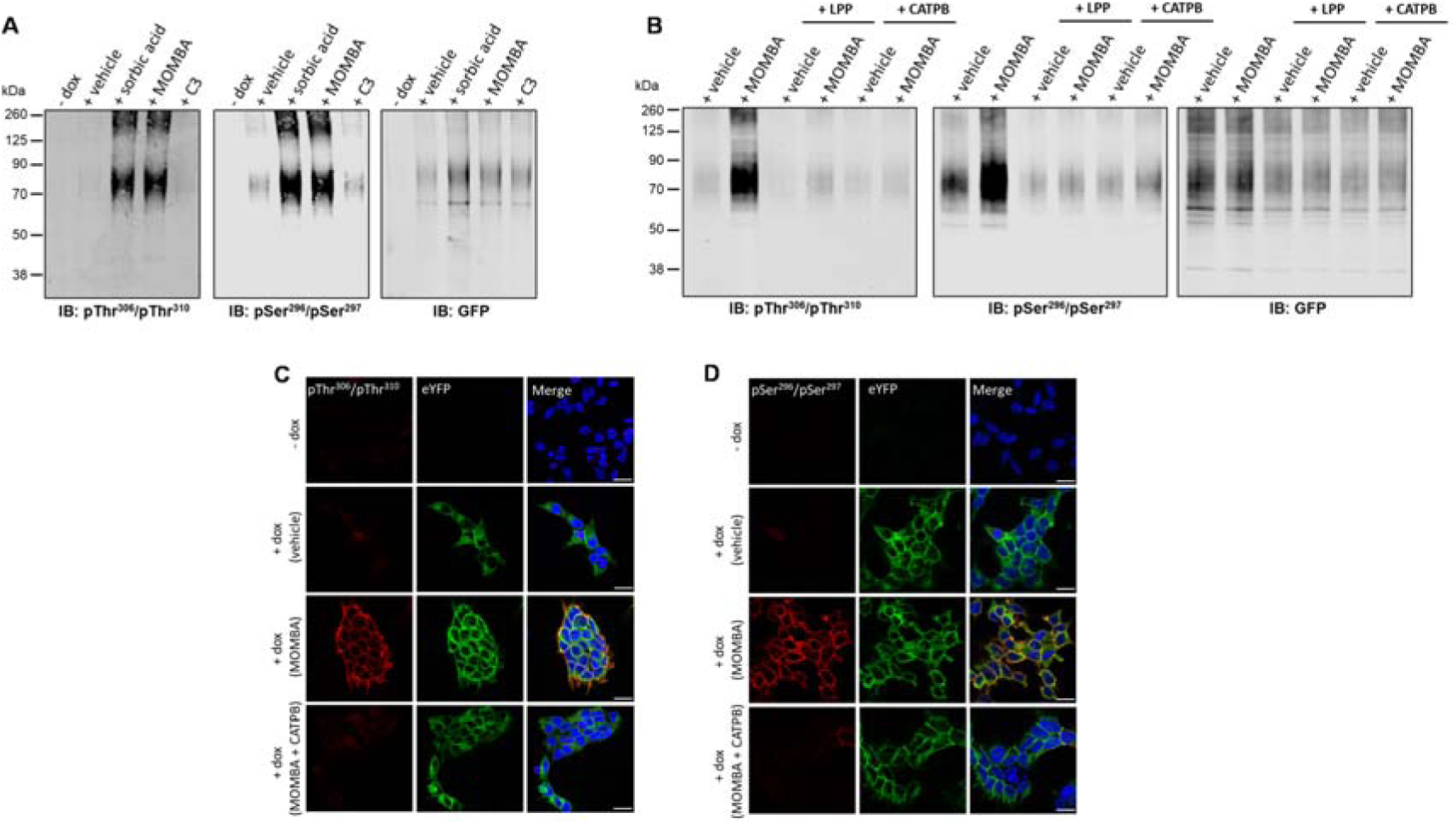
Agonist-induced detection of hFFA2-DREADD with putative pSer^296^ /pSer^297^ and pThr^306^ /pThr^310^ antisera reflects receptor activation, receptor phosphorylation and can be detected *in situ*. The ability of the pSer^296^ /pSer^297^, pThr^306^ /pThr^310^ hFFA2 and, as a control GFP, antisera to identify hFFA2-DREADD-eYFP after induction to express the receptor construct and then treatment of cells with vehicle, MOMBA, sorbic acid (each 100 μM) or propionate (C3) (2 mM) is shown. In the ‘**-dox**’ lanes receptor expression was not induced. **B**. as in **A**. except that after cell treatment with vehicle or MOMBA, immune-enriched samples were treated with Lambda Protein Phosphatase (LPP) or, rather than treatment with MOMBA, cells were treated with a combination of MOMBA and the hFFA2 inverse agonist CATPB (10 μM, 30 min pre-treatment). **C, D**. Cells harboring hFFA2-DREADD-eYFP and grown on glass coverslips were either untreated (- **dox**) or induced to express hFFA2-DREADD-eYFP. The induced cell samples were then exposed to vehicle, MOMBA (100 μM) or a combination of MOMBA (100 μM) and CATPB (10 μM) for 5 min. Fixed cells were then treated with anti-pThr^306^ /pThr^310^ (**C**) or anti-pSer^296^ /pSer^297^ (**D**) (**red**, Alexa Fluor 647) or imaged to detect eYFP (**green**). DAPI was added to detect DNA and highlight cell nuclei (**blue**). Scale bar = 20 μm.

To further confirm that the effects of MOMBA were due to direct activation of hFFA2-DREADD-eYFP, we assessed whether the hFFA2 receptor orthosteric antagonist/inverse agonist CATPB ((*S*)-3-(2-(3-chlorophenyl)acetamido)-4-(4- (trifluoromethyl)phenyl)butanoic acid) (32,33) would be able to prevent the effects of MOMBA. Indeed, pre-addition of CATPB (10 μM) to cells induced to express hFFA2- DREADD-eYFP prevented the effects of MOMBA on receptor recognition by both the putative pSer^296^/pSer^297^ and the pThr^306^/pThr^310^ antisera (**Figure 3B**). Additionally, pre-addition of CATPB lowered the extent of basal detection of the receptor by the pSer^296^/pSer^297^ antiserum (**Figure 3B**). As CATPB possesses inverse agonist properties (32) this may indicate that at least part of the basal pSer^296^/pSer^297^ signal reflects agonist-independent, constitutive activity of hFFA2-DREADD-eYFP. Once more, parallel immunoblots using anti-GFP confirmed that there were similar levels of the receptor construct in vehicle and MOMBA +/- CATPB-treated samples (**Figure 3B**). To conclude the initial characterization of the antisera we confirmed that both the putative pSer^296^/pSer^297^and pThr^306^/pThr^310^ antisera were detecting phosphorylated states of hFFA2-DREADD-eYFP. To do so, after exposure of cells induced to express hFFA2-DREADD-eYFP to either vehicle or MOMBA samples were treated with Lambda Protein Phosphatase (LPP) prior to SDS-PAGE and immunoblotting. This treatment is anticipated to remove phosphate from proteins in a site-agnostic manner. Now, MOMBA-induced detection of the receptor by both pSer^296^/pSer^297^ and pThr^306^/pThr^310^ antisera was eliminated (**Figure 3B**), as was most of the agonist-independent detection by pSer^296^/pSer^297^ (**Figure 3B**).

### MOMBA-induced phosphorylation of hFFA2-DREADD-eYFP is also detected in immunocytochemical studies

The pThr^306^/pThr^310^ antiserum was also effective in detecting MOMBA-induced post-activation states of hFFA2-DREADD-eYFP in immunocytochemistry studies. When induced in Flp-In T-REx 293 cells the presence of the receptor construct could be observed via the eYFP tag with or without exposure to MOMBA (**Figure 3C**). However, also in this setting, the anti-pThr^306^/pThr^310^ antiserum was only able to identify the receptor after treatment with MOMBA (**Figure 3C**) and, once again, pre-addition of CATPB prevented the effect of MOMBA (**Figure 3C**) without affecting direct identification and imaging of the receptor via the eYFP tag. Similar studies were performed with the hFFA2 pSer^296^/pSer^297^ antiserum with very similar outcomes. In this setting identification of hFFA2 DREADD-eYFP by the pSer^296^/pSer^297^ antiserum was almost entirely dependent on exposure to MOMBA (**Figure 3D**) and this was also prevented by pre-treatment with CATPB (**Figure 3D**).

### Phospho-specific antisera differentially identify hFFA2-DREADD in mouse tissues

To assess physiological roles of FFA2 in mouse tissues without potential confounding effects of either activation of the related SCFA-receptor FFA3 or non-receptor mediated effects of SCFAs we recently developed a hFFA2-DREADD knock-in transgenic mouse line (29). Here, mouse FFA2 is replaced by hFFA2-DREADD with, in addition, an appended C-terminal anti-HA epitope tag sequence to allow effective identification of cells expressing the receptor construct (20, 28).

#### White adipose tissue

FFA2 is known to be expressed in white adipose tissue and we previously used these hFFA2-DREADD-HA mice to define the role of this receptor as an anti-lipolytic regulator (20). To explore the expression and regulation of hFFA2-DREADD-HA more fully we took advantage of the appended HA tag to immunoprecipitate the receptor from white adipose tissue taken from hFFA2-DREADD-HA mice. As a control, equivalent pull-down studies were performed with tissue from ‘CRE-MINUS’ animals (20, 28). These harbor hFFA2-DREADD-HA at the same genetic locus but expression has not been induced and hence they lack protein corresponding to either hFFA2-DREADD-HA or mouse FFA2 (20). In all of these studies, and those using other mouse-derived tissues (see later), as well as an anti-protease cocktail, we included the phosphatase inhibitor cocktail PhosSTOP, as described previously (34), to prevent potential dephosphorylation of hFFA2-DREADD-HA and other proteins. Following SDS-PAGE of such immune-precipitates from hFFA2- DREADD-HA mice immunoblotting with an HA-antibody resulted in detection of hFFA2- DREADD-HA predominantly as an approximately 45 kDa species, with lower levels of a 40 kDa form (**Figure 4A**). These both clearly corresponded to forms of hFFA2-DREADD-HA as such immunoreactive polypeptides were lacking in pull-downs from white adipose tissue from CRE-MINUS mice (**Figure 4A**). Moreover, parallel immunoblotting with an antiserum against the non-post-translationally modified C-terminal tail sequence of human FFA2 identified the same polypeptides as the HA-antibody and, once more, this was only in tissue from the hFFA2-DREADD-HA and not CRE-MINUS animals (**Figure 4A**). Notably, immunoblotting of such samples with the anti-pSer^296^/pSer^297^ antiserum also detected the same polypeptides. However, unlike the hFFA2 C-terminal tail antiserum, although the pSer^296^/pSer^297^ antiserum did identify hFFA2-DREADD-HA without prior addition of a ligand, pre-addition of MOMBA increased immune-detection of the receptor (**Figure 4A**). This is consistent with the ligand promoting quantitatively greater levels of phosphorylation of these residues (**Figure 4B**). Moreover, pre-addition of the hFFA2 antagonist/inverse agonist CATPB not only prevented the MOMBA-induced enhanced detection of the receptor protein by the pSer^296^/pSer^297^ antiserum but showed a trend to reduce this below basal levels (**Figure 4A**) although this did not reach statistical significance (**Figure 4B**). In contrast, the anti-pThr^306^/pThr^310^ antiserum failed to detect immunoprecipitated hFFA2-DREADD-HA from white adipose tissue in either the basal state or post-addition of MOMBA (**Figure 4A, 4B**). This indicates that these sites are not and do not become phosphorylated in this tissue.

**Figure 4.**
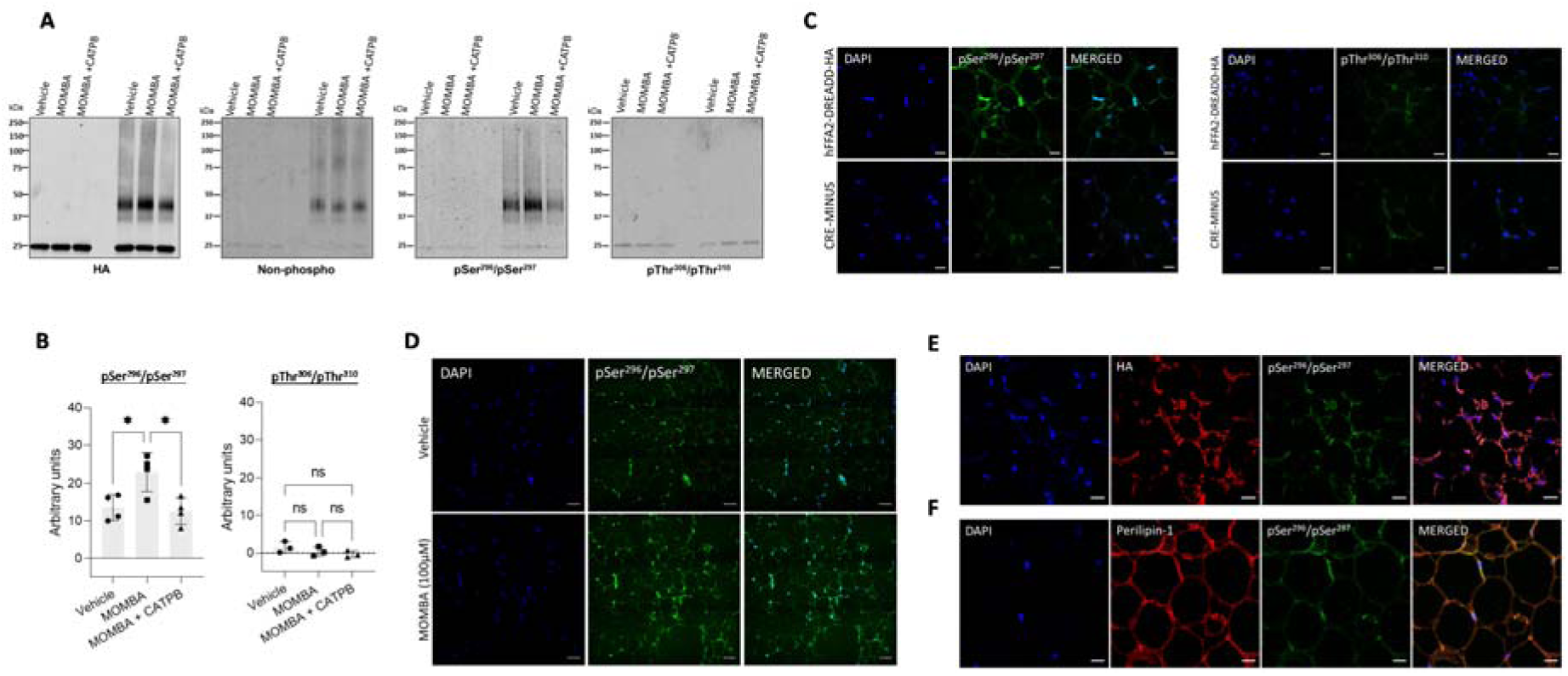
In white adipose tissue residues Ser^296^ /Ser^297^ of hFFA2-DREADD-HA but not Thr^306^ /Thr^310^ become phosphorylated in response to MOMBA. White adipose tissue dissected from hFFA2-DREADD-HA and CRE-MINUS mice was treated with either vehicle, 100 μM MOMBA or 100 μM MOMBA + 10 μM CATPB **A.** Lysates were prepared and solubilized. hFFA2-DREADD-HA was immunoprecipitated using an anti-HA monoclonal antibody and following SDS-PAGE immunoblotted to detect HA, non-phosphorylated hFFA2-DREADD-HA, and pSer^296^ /pSer^297^ or pThr^306^ /pThr^310^ hFFA2-DREADD-HA. A representative experiment is shown. **B.** Quantification of pSer^296^ /pSer^297^ (**left**) and pThr^306^ /pThr^310^ immunoblots (**right**) phosphorylation (means +/- S.E.M.) in experiments using tissue from different mice, * p < 0.05, ns: not significant. **C.** Tissue samples from hFFA2-DREADD-HA (**top panel**) and CRE-MINUS (**bottom panel**) mice that were treated with MOMBA were immunostained to detect pSer^296^ /pSer^297^ (**left panels**), pThr^306^ /pThr^310^ (**right panels**) and counterstained with DAPI (**blue**). Scale bars = 20 μm. **D.** Comparison of pSer^296^ /pSer^297^ staining of samples from hFFA2-DREADD-HA expressing mice vehicle treated (**top panels**) or treated with MOMBA (**bottom panels**) (scale bar = 50 μm). **E, F**. Tissue sections from hFFA2-DREADD-HA-expressing mice immunostained with pSer^296^ /pSer^297^ (**green**) and anti-HA (**red**) (**E**) to detect the receptor expression or **F** with anti-perilipin-1 (**red**) to identify adipocytes. Merged images are shown to the right. Scale bars = 20 μm.

Immunohistochemical studies performed on fixed adipose tissue from hFFA2- DREADD-HA expressing mice that had been pre-exposed to MOMBA were consistent with the immunoblotting studies. Clear identification of pSer^296^/pSer^297^ hFFA2-DREADD-HA was detected and the specificity of this was confirmed by the absence of staining in equivalent samples from CRE-MINUS mice (**Figure 4C**). Once more, no specific staining was observed for the anti-pThr^306^/pThr^310^ antiserum (**Figure 4C**). Clear detection of pSer^296^/pSer^297^ immunostaining was observed in adipose tissue from hFFA2-DREADD-HA mice without exposure to MOMBA but once more the intensity of staining was markedly increased after MOMBA treatment (**Figure 4D**). Parallel anti-HA staining showed strong co- localization with the pSer^296^/pSer^297^ antiserum (**Figure 4E**), and clear co-localization of anti-pSer^296^/pSer^297^ immunostaining with that for perilipin-1 (**Figure 4F**) confirmed the presence of pSer^296^/pSer^297^ hFFA2-DREADD-HA directly on adipocytes.

#### Peyer’s patch immune cells

As the apparent lack of anti-pThr^306^/pThr^310^ immunostaining after treatment with MOMBA in white adipocytes was distinct from the outcomes observed in Flp-In T-REx 293 cells we turned to a second source of tissue in which FFA2 expression is abundant. Peyer’s patches act as immune sensors of the gut (35). Anti-HA staining showed extensive and high-level expression of hFFA2-DREADD-HA in cells within these structures (**Figure 5A**). Co-staining for CD11c indicated many of the hFFA2-DREADD-HA positive cells correspond to dendritic cells, monocytes and/or macrophages (**Figure 5B**). Co-staining to detect the presence of the nuclear transcription factor RoRγt also indicated, as shown previously at the mRNA level for mouse FFA2 (36), that hFFA2-DREADD-HA is well expressed by type-III innate lymphoid cells (**Figure 5C**). HA pull-down immune-captured hFFA2-DREADD-HA from Peyer’s patches and associated mesenteric lymph nodes of hFFA2-DREADD-HA expressing mice (**Figure 5D**). Anti-HA immunoblotting of such material indicated a more diffuse pattern of immunostaining than observed from white adipose tissue (**Figure 5D**). This may reflect a more complex pattern of post-translational modifications, including differing extents of N-glycosylation (see later) than observed in white adipose tissue (compare **Figure 5D** with **Figure 4A**). Once more, however, this range of HA-detected polypeptides did indeed all represent forms of hFFA2-DREADD-HA as they were completely absent from HA-immunocapture conducted in equivalent tissue from the CRE-MINUS animals (**Figure 5D**). Parallel immunoblots of such samples with the anti-pThr^306^/pThr^310^ antiserum now indicated that, in contrast to adipocytes, hFFA2-DREADD-HA became phosphorylated on these residues in a MOMBA-dependent manner, with no detection of phosphorylation of these residues without agonist treatment (**Figure 5D**).

**Figure 5.**
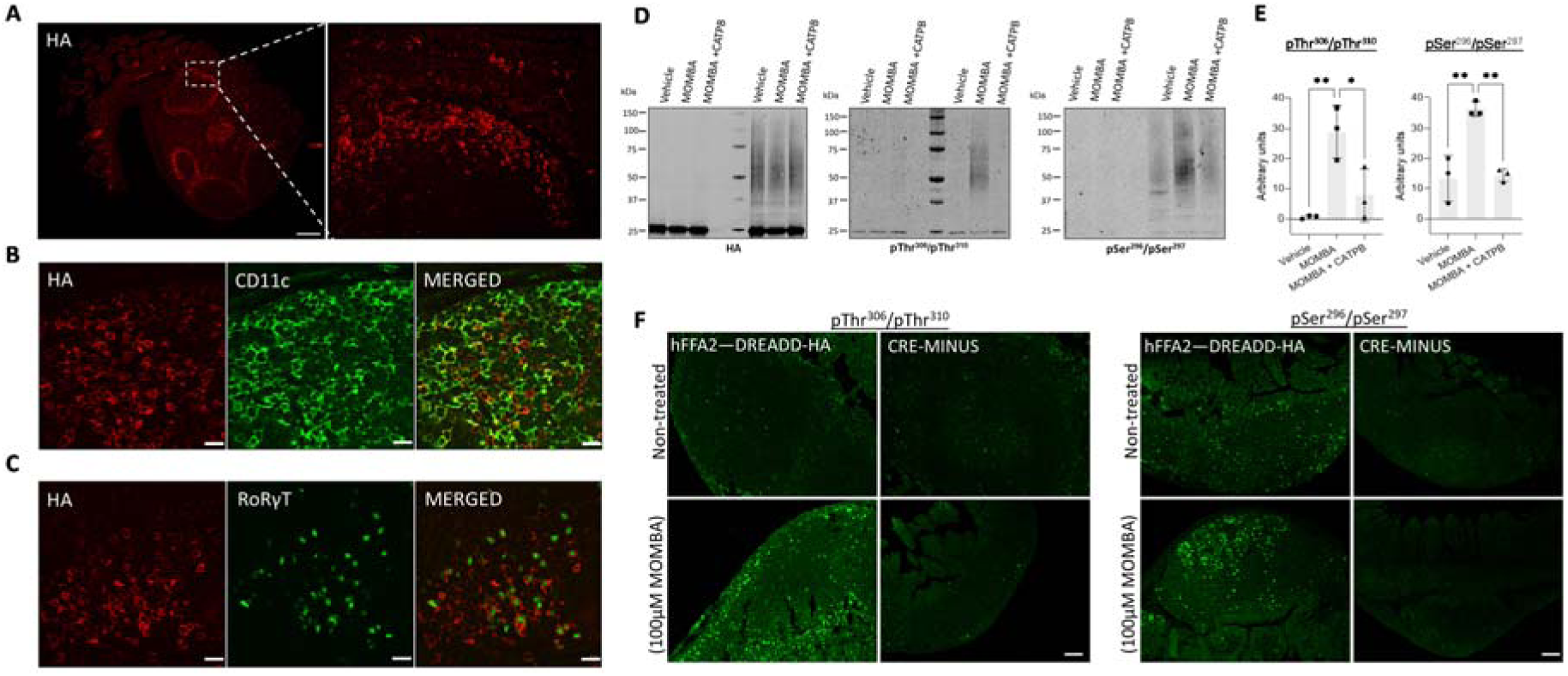
hFFA2-DREADD-HA becomes phosphorylated at Thr^306^ /Thr^310^ in addition to Ser^296^ /Ser^297^ in immune cells from Peyer’s patches. Peyer’s patches isolated from hFFA2-DREADD-HA expressing mice were immunostained with anti-HA (**red**) to detect receptor expression. Images were acquired with ×20 (**left panel**) and ×63 (**right panel**) objectives (scale bar = 200 μm) (**A**). Tissue sections were counterstained with **B.** anti-CD11c as a marker of dendritic cells, monocytes and/or macrophages or, **C.** ROR©T to detect type-III innate lymphoid cells (scale bar = 20 μm). Isolated Peyer’s patches and mesenteric lymph nodes from CRE-MINUS and hFFA2- DREADD-HA mice were exposed to either vehicle, 100 μM MOMBA or 100 μM MOMBA + 10 μM CATPB. **D.** Following lysate preparation, immunoprecipitation and SDS-PAGE samples were probed to detect HA, pThr^306^ /pThr^310^ or pSer^296^ /pSer^297^. **E.** Quantification of pThr^306^ /pThr^310^ (**left**) and pSer^296^ /pSer^297^ immunoblots (**right**) hFFA2- DREADD-HA (means +/- S.E.M.), *p < 0.05, **p < 0.01. **F.** Treated tissue sections were also used in immunohistochemical studies, employing either pThr^306^ /pThr^310^ (**left panels**) or pSer^296^ /pSer^297^ (**right panels**) (scale bars = 100 μm).

Moreover, this effect of MOMBA was clearly mediated by the hFFA2-DREADD-HA receptor because anti-pThr^306^/pThr^310^ recognition was entirely lacking when MOMBA was added after the addition of the hFFA2 antagonist/inverse agonist CATPB (**Figures 5D, 5E**) that blocks this receptor with high affinity (32,33). Similar outcomes were obtained in this tissue with the anti-pSer^296^/pSer^297^ antiserum. Basal detection of the immune-captured receptor was low and this was increased markedly by pre-treatment with MOMBA (**Figures 5D, 5E**). As for the anti-pThr^306^/pThr^310^ antiserum the effect of MOMBA on recognition of the receptor was not observed when cells were exposed to CATPB in addition to MOMBA (**Figures 5D, 5E**). Immunostaining of fixed Peyer’s patch tissue with the anti-pThr^306^/pThr^310^ antiserum confirmed the marked agonist-dependence of recognition of hFFA2-DREADD-HA in such cells (**Figure 5F**). We detected a low level of immunostaining with this antiserum in Peyer’s patch tissue in the absence of addition of MOMBA but, as we detected a similar level of staining in tissue from CRE-MINUS animals, in both in the presence and absence of MOMBA, this may represent a small level of non-specific reactivity (**Figure 5F**). In this tissue, immunostaining with the pSer^296^/pSer^297^ antiserum indicated a degree of agonist-independent phosphorylation of these sites as this was not observed in tissue from CRE-MINUS mice, and a further increase in staining following pre-treatment with MOMBA was evident (**Figure 5F**). Higher level magnification allowed detailed mapping of the location of pThr^306^/pThr^310^ hFFA2-DREADD-HA (**Supplementary Figure 1**).

Thus, although basal and agonist-regulated phosphorylation of Ser^296^/Ser^297^ hFFA2- DREADD-HA was evident in both white adipocytes and Peyer’s patch immune cells, activation-induced Thr^306^/Thr^310^ phosphorylation is observed in immune cells but not in white adipose tissue.

#### Lower gut enteroendocrine cells

We have previously used anti-HA and both anti-GLP-1 and anti-PYY antisera to illustrate the expression of hFFA2-DREADD-HA in GLP-1 and PYY-positive entero-endocrine cells of the colon of these mice and shown that activation of this receptor enhances release of both GLP-1 (20) and PYY (28). To extend this we added vehicle or MOMBA to preparations of colonic epithelia from these mice. Subsequent immunostaining with the anti-pThr^306^/pThr^310^ antiserum identified a limited number of widely dispersed cells in MOMBA-treated tissue but minimal staining of the vehicle-treated controls (**Figure 6A, 6B**). This highlighted groups of spatially-scattered cells in which hFFA2-DREADD-HA became phosphorylated on these residues in an agonist-dependent manner (**Figure 6B**). As an additional specificity control similar experiments were performed with the anti-pThr^306^/pThr^310^ antiserum on tissue from the CRE-MINUS mice. These failed to identify equivalent cells (**Figure 6A, 6B**). Despite the limited cell expression pattern of anti-HA detected previously in colonic tissue (20) we were again able to specifically capture hFFA2-DREADD-HA via HA-pulldown (**Figure 6C**). Here, although challenging to detect, we were able to record specific agonist-mediated immuno-recognition of polypeptides of the appropriate molecular mass with both the anti-pThr^306^/pThr^310^ and anti-pSer^296^/pSer^297^ antisera that were again absent following HA pull-downs conducted in tissue from CRE-MINUS mice (**Figure 6C**). However, compared to the intensity of immunodetection of either anti-pThr^306^/pThr^310^ and/or anti-pSer^296^/pSer^297^ in white adipose tissue or Peyer’s patch immune cells compared to the extent of anti-HA pulldown, such detection was modest and potentially indicted only limited phosphorylation of these sites in such colonic epithelial cells, even after exposure to MOMBA (**Figure 6C**).

**Figure 6.**
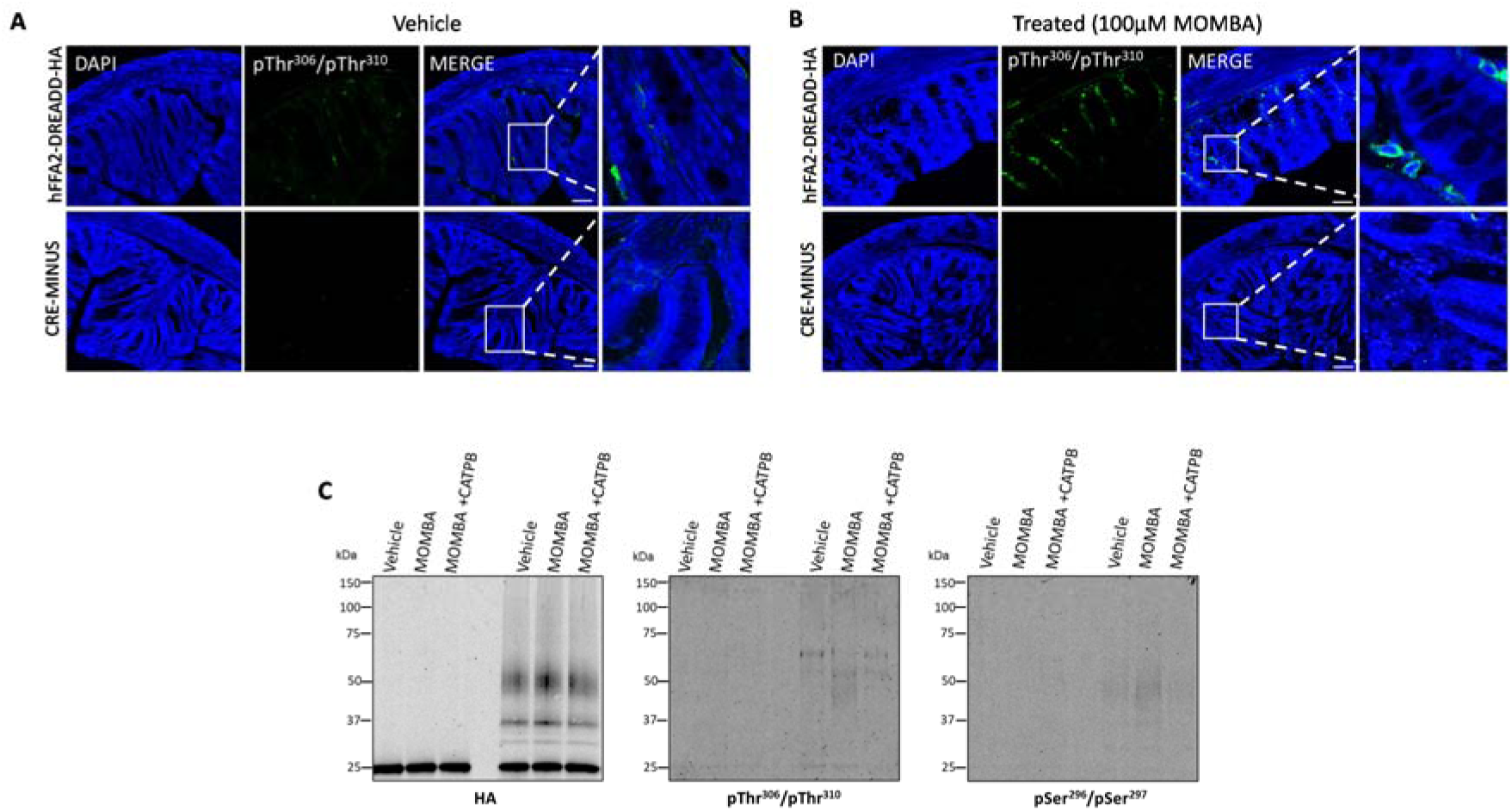
MOMBA promotes limited phosphorylation of both Ser^296^ /Ser^297^ and Thr^306^ /Thr^310^ in hFFA2-DREADD-HA in lower gut enteroendocrine cells A. B., Colonic tissue isolated from hFFA2-DREADD-HA **(top panels)** or CRE-MINUS **(bottom panels)** mice treated with either vehicle **(A)** or 100 μM MOMBA **(B)**. Following fixation tissue sections were immunostained with pThr^306^ /pThr^310^ and counterstained with DAPI. (Scale bar = 100 μm). In the merged images, the box is expanded in the right-hand panels. **C**. Lysates prepared from tissue samples treated as noted were analysed by probing immunoblots with anti-HA, anti-pThr^306^ /pThr^310^ or anti-pSer^296^ /pSer^297^. Representative examples are shown.

### Wild type hFFA2 shows marked similarities in regulated phosphorylation to hFFA2- DREADD

Whilst the DREADD receptor strategy offers unique control over ligand activation of a GPCR and limits potential activation in tissues by circulating endogenous ligands, it is important to assess if similar outcomes are obtained when using the corresponding wild type receptor. To do so we next employed Flp-In T-REx 293 cells that allowed doxycycline-induced expression of wild type hFFA2-eYFP. As the wild type and DREADD forms of hFFA2 are identical within their C-terminal regions it was anticipated that the anti-pSer^296^/pSer^297^ and anti-pThr^306^/pThr^310^ antisera would be able to detect phosphorylation of these residues in wild type hFFA2-eYFP, as noted for hFFA2-DREADD-eYFP, but instead regulated by SCFAs such as propionate (C3) rather than by MOMBA. Immunoblots using lysates from these cells with either of these antisera showed predominant identification of a group of polypeptides migrating with apparent mass in the region of 60 kDa after treatment of cells induced to express hFFA2-eYFP with C3 (2 mM, 5 min) (**Figure 7A**). Recognition in this setting of hFFA2-eYFP by anti-pThr^306^/pThr^310^ was almost completely dependent on addition of C3 whilst, although also markedly enhanced by addition of C3, there was some level of identification of hFFA2-eYFP by anti-pSer^296^/pSer^297^ without addition of C3 (**Figure 7A**). Parallel immunoblotting with anti-GFP confirmed the presence of similar levels of hFFA2-eYFP both with and without treatment with C3 (**Figure 7A**) and confirmed in addition that expression of the receptor construct was lacking if cells harbouring hFFA2- eYFP had not been induced with doxycycline (**Figure 7A**). As with the effect of MOMBA on the hFFA2-DREADD-eYFP construct, in studies using the anti-pThr^306^/pThr^310^ antiserum, C3-induced recognition of the hFFA2-eYFP receptor was lacking in the co-presence of the hFFA2 antagonist CATPB (**Figure 7B**). In addition, treatment of C3-exposed samples to LPP before SDS-PAGE also prevented subsequent hFFA2-eYFP identification by the anti-pThr^306^/pThr^310^ antiserum (**Figure 7B**), confirming this to reflect phosphorylation of the target.

**Figure 7.**
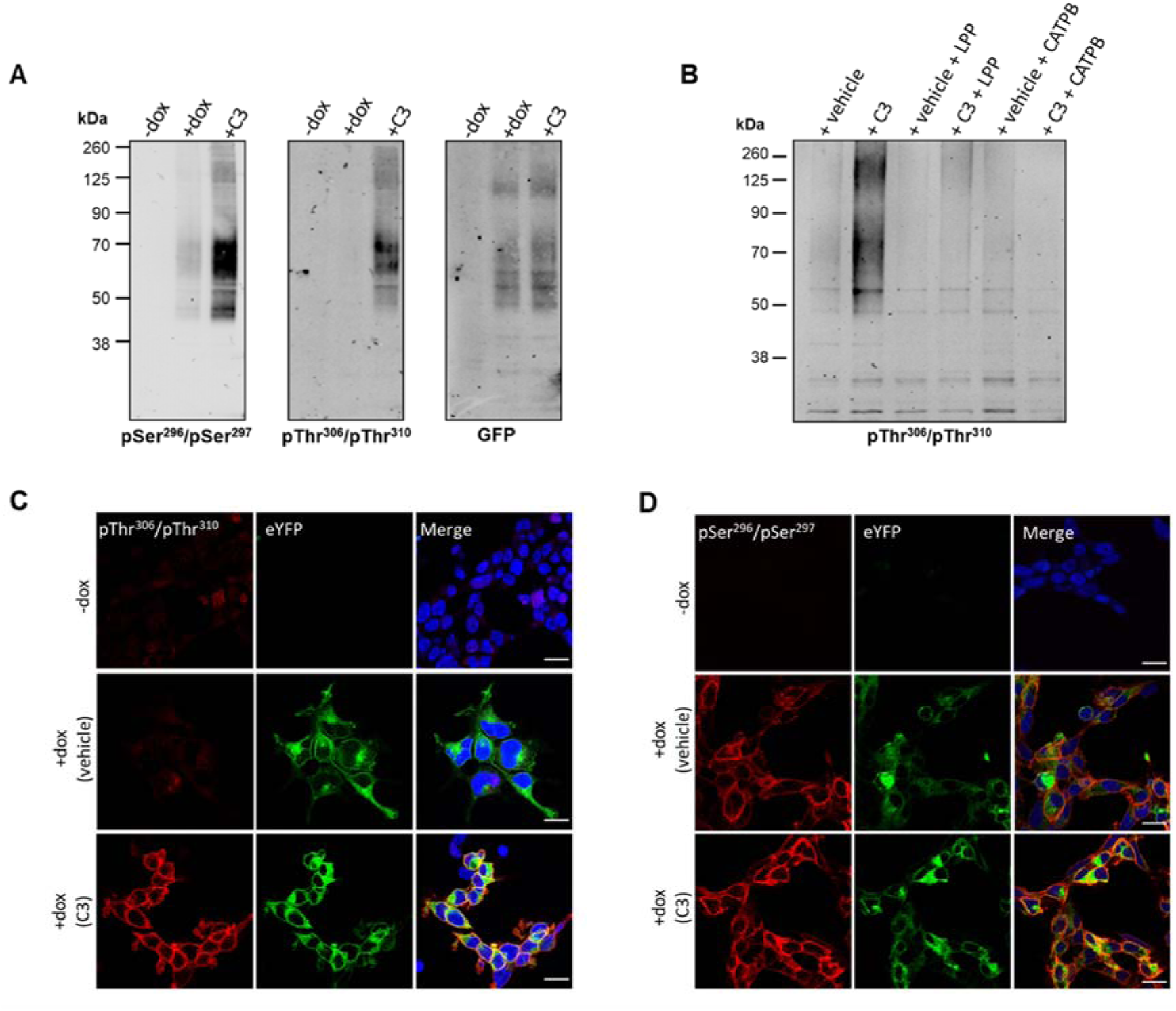
Propionate regulates phosphorylation of hFFA2-eYFP: *In vitro* studies. Flp-In T-REx 293 cells habouring hFFA2-eYFP were induced to express the receptor construct (**+ dox**) or not (**- dox**) and the induced cells were then treated with propionate (C3, 2 mM, 5 min) or vehicle. **A**. Cell lysates were resolved by SDS-PAGE and then immunoblotted with anti-pSer ^296^ /pSer ^297^ hFFA2, anti-pThr ^306^ /pThr ^310^ hFFA2, or anti-GFP. **B**. Cells induced to express hFFA2-eYFP were treated with C3 (2 mM, 5 min) or vehicle. Where noted cells were pre-treated with the hFFA2 antagonist CATPB (10 μM, 20 min before agonist addition). Lysates were then prepared and, where indicated, treated with Lambda Protein Phosphatase (LPP). Following SDS-PAGE the samples were immunoblotted with anti-pThr ^306^ /pThr ^310^ hFFA2. **C**., **D**. Cells were doxycycline induced (+ dox) or not (-dox) and prepared for immunocytochemistry after treatment with C3 or vehicle and exposed to anti-pThr ^306^ /pThr ^310^ hFFA2 (**C**) or anti-pSer ^296^ /pSer ^297^ hFFA2 (**D**) (**red**) whilst direct imaging detected the presence of hFFA2-eYFP (**green**). Merged images (**right hand panels**) were also stained with DAPI (**blue**) to identify cell nuclei. Scale bars = 20 μm.

### C3-induced phosphorylation of hFFA2-eYFP is also detected in immunocytochemical studies

As an alternate means to assess the phosphorylation status of hFFA2-eYFP in Flp-In T-REx 293 cells we also performed immunocytochemical studies after cells induced to express the receptor construct had been exposed to either C3 (2 mM) or vehicle. Imaging the presence of eYFP confirmed receptor expression in each case (**Figures 7C, 7D**) and, in addition, lack of the receptor in cells that had not been exposed to doxycycline. The anti-pThr^306^/pThr^310^ antiserum was only able to identify hFFA2-eYFP after exposure to C3 in this context (**Figure 7C**), whereas for this construct the anti-pSer^296^/pSer^297^ antiserum identified the receptor in both vehicle and C3-treated cells (**Figure 7D**).

### Studies in tissues from transgenic mice expressing hFFA2-HA

To expand such comparisons into native tissues we generated a further transgenic knock-in mouse line. Here we replaced mFFA2 with hFFA2-HA, again with expression of the transgene being controlled in a CRE-recombinase dependent manner. To assess the relative expression of hFFA2-HA in this line compared to hFFA2-DREADD-HA in equivalent tissues of the hFFA2-DREADD-HA knock-in mice we isolated both white adipocytes and colonic epithelia from homozygous animals of each line and immunoprecipitated the corresponding receptors using anti-HA. Immunoblotting of such HA-immunoprecipitates showed similar levels of the corresponding receptor in each and that the molecular mass of hFFA2-HA was equivalent to hFFA2-DREADD-HA **(Supplementary Figure 2).**

In immunocytochemical studies using Peyer’s patches isolated from the hFFA2-HA expressing mice addition of C3 (10 mM, 20 min) produced a marked increase in detection of hFFA2-HA by the anti-pThr^306^/pThr^310^ antiserum compared to vehicle-treated samples (**Figure 8A**). By contrast, such an effect of C3 was lacking in equivalent tissue isolated from equivalent CRE-MINUS comparator mice (**Figure 8A**). Basal identification of anti-pSer^296^/pSer^297^ staining was significant (**Figure 8B**) and little altered by treatment with C3 (**Figure 8B**). Such basal phosphorylation of Ser^296^/Ser^297^ did appear to be specific however, as this was lacking in tissue from the CRE-MINUS mice (**Figure 8B**). To extend these observations we again used anti-HA to immunoprecipitate hFFA2-HA from Peyer’s patches from hFFA2-HA expressing and CRE-MINUS animals after treatment with vehicle, C3, or C3 + CATPB. As we observed in samples from hFFA2-DREADD-HA expressing mice, subsequent to SDS-PAGE the receptor isolated from such cells migrated as a smear of anti-HA immunoreactivity with apparent molecular masses ranging from some 50-70 kDa (**Figure 8C**). However, this complex mix clearly corresponded to forms of hFFA2-HA as such immunoreactivity was once more lacking in HA-pulldowns of tissue from CRE-MINUS mice (**Figure 8C**). Immunoblotting such samples with the anti-pThr^306^/pThr^310^ antiserum showed clear phosphorylation of pThr^306^/pThr^310^ in response to C3 (**Figure 8C**) that increased from essentially undetectable levels in samples of vehicle-treated cells (**Figure 8D**).

**Figure 8.**
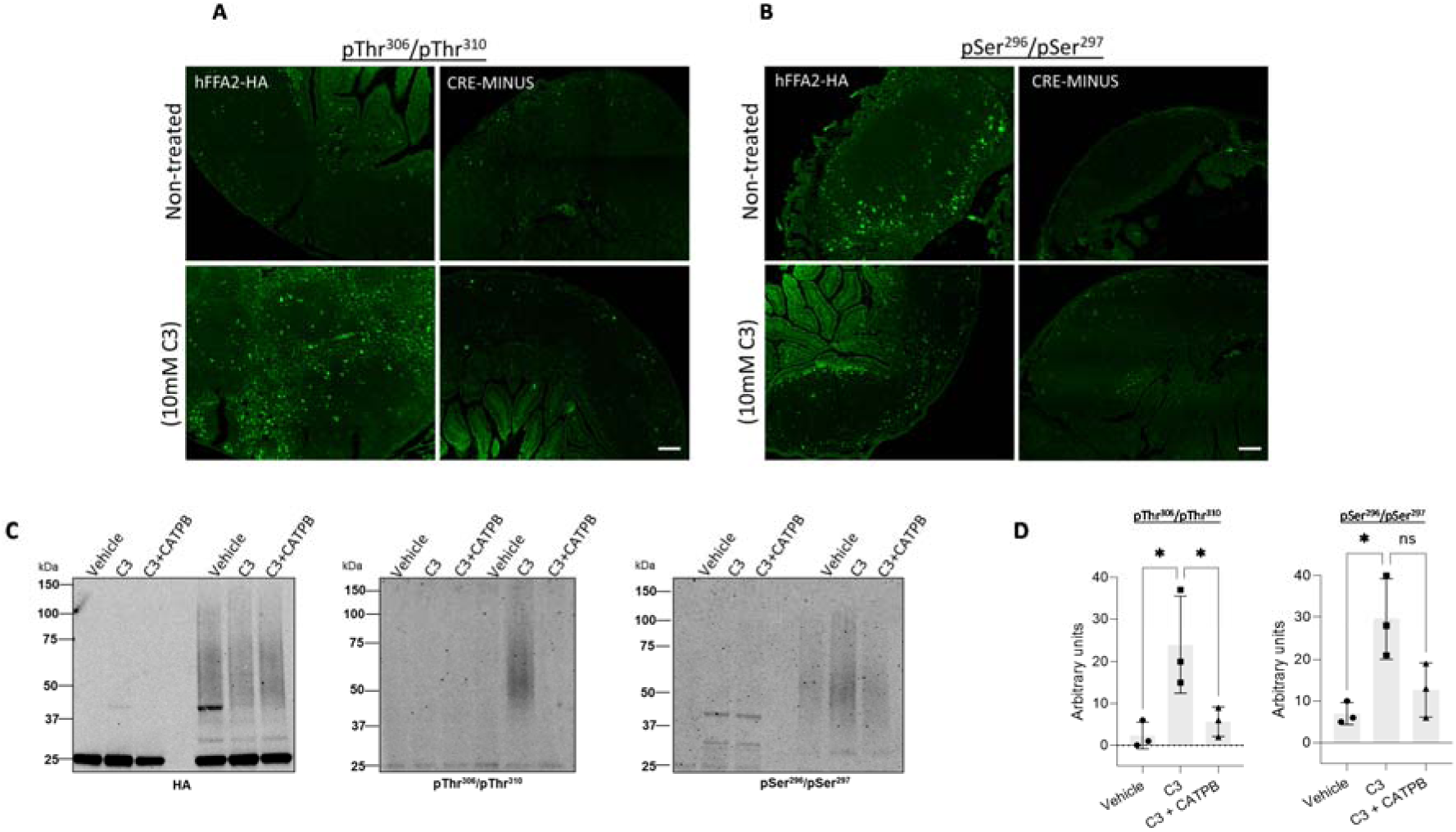
C3-induces phosphorylation of both pSer^296^/pSer^297^ and pThr^306^/pThr^310^ in Peyer’s patches from hFFA2-HA expressing mice. Isolated Peyer’s patches and mesenteric lymph nodes from hFFA2-HA and the corresponding CRE-MINUS mice were exposed to either vehicle, or 10 mM C3 for 20 minutes. Tissue sections were used in immunohistochemical studies, employing either anti-pThr ^306^ /pThr ^310^ (**A**) or anti-pSer ^296^ /pSer ^297^ (**B**) (scale bars = 100 μm). **C.** Lysates from Peyer’s patches isolated from hFFA2-HA expressing mice, or the corresponding CRE-MINUS mice, that had been treated with vehicle, C3 (10 mM, 20 min), or C3 + CATPB (10 μM, 30 minutes before agonist) were immunoprecipitated with anti-HA as for the hFFA2-DREADD-HA expressing mice in Figure 5. Subsequent to SDS-PAGE samples such were probed to detect HA (**C, left**), anti-pThr ^306^ /pThr ^310^ (**C, centre**) or anti-pSer ^296^ /pSer ^297^ (**C, right)**. hFFA2-HA was detected as a broad smear of protein(s) with Mr centred close to 55 kDa. **D.** Quantification of pThr ^306^ /pThr ^310^ (**left**) and pSer ^296^ /pSer ^297^ immunoblots (**right**) phosphorylation in experiments using tissue from three different mice (means +/- S.E.M.), * p < 0.05, ns: not significant.

Like cells from hFFA2-DREADD-HA expressing Peyer’s patches the effect of C3 was clearly produced directly at the hFFA2-HA receptor because pre-treatment with CATPB entirely blocked the effect of C3 (**Figures 8C, 8D**). The anti-pSer^296^/pSer^297^ antiserum also showed enhanced detection of hFFA2-HA in response to treatment with C3 in this setting that was also prevented by pre-treatment with CATPB (**Figures 8C, 8D**) but this appeared to be less pronounced than in the studies employing anti- pThr^306^/pThr^310^ hFFA2.

## Discussion

The ability to detect activated GPCRs in native tissues can provide insights into how such receptors are regulated and, in drug-discovery and target validation efforts, provide valuable information on target engagement. For most GPCRs, as well as interaction with members of the family of heterotrimeric G proteins, upon agonist occupancy this also results in rapid phosphorylation of various serine and threonine residues, usually within the third intracellular loop and/or the C-terminal tail. Often mediated by members of the GRK family of kinases such phosphorylation allows higher affinity interactions with arrestin proteins. Classically this is linked to desensitization of receptor-mediated second messenger regulation, as binding of an arrestin usually occludes the G protein binding site. However, now and for a considerable period, it has been established that arrestin interactions can promote distinct signalling functions and that these might be dependent on the detailed architecture of the GPCR-arrestin complex (37, 38). This can vary with the pattern of agonist-induced GPCR phosphorylation (39, 40). This concept of phosphorylation-barcoding (37, 39-40) has been enhanced greatly by both improved approaches to detect such sites via advances in mass spectrometry (41,42) and the development and use of phosphorylation-site specific antisera (31, 40, 43-44). Here, guided by combinations of mass spectrometry-identified sites of basal and agonist-regulated phosphorylation of a chemogenetically-modified form of the SCFA responsive receptor FFA2, and informatic predictions of sites that might be or become phosphorylated in an activation-dependent manner, we generated three distinct sets of antisera. For one of these, potentially targeting pSer^324^/pSer^325^ hFFA2-DREADD, we had no direct evidence from mass spectrometry performed on the receptor enriched from a HEK293- derived cell line expressing hFFA2-DREADD-eYFP of phosphorylation of these sites.

Moreover, we failed to find evidence to support such phosphorylation, using the generated antiserum, in either the cell line or the mouse tissues we studied. As we will discuss for the other antisera, this does not however explicitly exclude that these sites might be phosphorylated in other tissues or settings. For the anti-pThr^306^/pThr^310^ hFFA2-DREADD antiserum we also did not find direct mass spectrometry support for phosphorylation of these residues in the HEK293-derived cell line. However, in both immunoblotting studies performed on lysates of these cells and in immunocytochemical studies, this antiserum was able to identify hFFA2-DREADD-eYFP in a sensitive and agonist-activation dependent manner. Moreover, experiments in which we denuded the receptor of phosphorylation by treatment with Lambda Protein Phosphatase (LPP) demonstrated that the antiserum was only able to identify the phosphorylated receptor and mutation to Ala of both Thr^306^ and Thr^310^ also eliminated detection of the receptor, confirming the sites of modification.

To assess aspects of potential differential bar-coding we required a second antiserum able to identify a different site(s) of phosphorylation in hFFA2-DREADD. Here, we did have direct mass spectrometry-derived evidence for basal phosphorylation of Ser^297^ and, in addition, agonist-dependent phosphorylation of Ser^296^. In the same way as discussed for the anti-pThr^306^/pThr^310^ antiserum we confirmed the specificity of this antiserum and showed in both immunoblotting studies and immunocytochemical studies that there was increased immunoreactivity of this antisera after cells expressing hFFA2-DREADD were exposed to an appropriate agonist.

Most significantly in the context of phosphorylation bar-coding we were able to show tissue-selective phosphorylation of Thr^306^ and Thr^310^. To do so we took advantage of a knock-in transgenic mouse line that we have characterized extensively (20, 28-29). In this line we replaced mouse FFA2 with hFFA2-DREADD. In addition, this line has an additional in-frame HA-epitope tag added to the C-terminus of the receptor. Both previously, and in the current studies, we have therefore detected the HA-tag to identify specific cells expressing the receptor. Herein, however, we built on this to identify the activated receptor and cells directly expressing the activated receptor. These studies provided direct evidence of tissue-specific patterns of agonist-induced FFA2 receptor bar-coding. In each of the three tissues we explored, white adipose tissue, immune cells in gut Peyer’s patches and in lower intestine enteroendocrine cells, we observed in immunocytochemical studies both basal and agonist-enhanced phosphorylation of Ser^296^/Ser^297^. By contrast, although we observed agonist- regulation of Thr^306^/Thr^310^ in both immune cells and enteroendocrine cells, these residues were not phosphorylated in white adipocytes. Hence, we provide clear evidence that the same agonist-GPCR pairing results in a different phosphorylation bar-code in adipocytes compared to Payer’s patch immune cells. In addition, the extent of phosphorylation of each of these sites was quantitatively much lower in colonic epithelial than either of the other tissues examined. As we have noted previously (34) successful outcomes of these experiments required the maintenance of phosphatase inhibitors throughout.

Importantly we also expanded these studies to a second transgenic knock-in mouse line in which we replaced mFFA2 with wild type hFFA2-HA. This was to assess whether the regulation of receptor phosphorylation we observed in tissues from the hFFA2-DREADD-HA expressing line might differ from the wild type receptor and, if so, whether this could possibly relate to the differences in the way the hFFA2-DREADD agonist MOMBA activates the receptor compared to the endogenous SCFAs at the wild type receptor. We considered this unlikely as we have previously shown indistinguishable characteristics of these two ligand-receptor pairs *in vitro* (20). Importantly, using HA-immunoprecipitation from equivalent tissues from these two lines we observed very similar levels of expression of hFFA2-DREADD-HA and hFFA2-HA in both white adipose tissue and colonic epithelial preparations and in each the receptor constructs migrated similarly in SDS-PAGE suggesting that co- and post-translational modifications were at least similar. Moreover, as the DREADD variant differs from wild type receptor only in two amino acids of the orthosteric ligand binding pocket, and the intracellular sections of the two forms are identical (20), we anticipated that the phospho-site specific antisera would recognise each in an equivalent manner. Overall, comparisons between tissues from the wild type hFFA2 and hFFA2- DREADD expressing animals were very similar. There was a hint that Ser^296^/Ser^297^ are more highly phosphorylated in the basal state when examining wild type hFFA2-HA mice but phosphorylation of Thr^306^/Thr^310^ was entirely agonist-dependent for both forms of the in both white adipose tissue and when using Flp-In T-REx 293 cells.

A further interesting feature, although not inherently linked to bar-coding, is that co-translational N-glycosylation, and potentially other post-translational modifications, of the hFFA2-DREADD-HA receptor (and probably of hFFA2-HA) was markedly different in these various tissues. As we were able to use the HA-tag to immunoprecipitate the receptor from different tissues, SDS-PAGE and immunoblotting studies showed that the predominant form of the receptor in colonic epithelium migrated as a substantially larger species than that isolated from white adipocytes, and this was also the case when the receptor was immune-enriched from Peyer’s patches and associated mesenteric lymph nodes (**Supplementary Figure 3**). De-glycosylation studies using an enzyme able to cleave N-linked carbohydrate confirmed these differences to reflect, at least in part, differential extents of N-glycosylation (**Supplemental Figure 3**).

The basis of the distinct bar-coding induced by MOMBA-activation of hFFA2- DREADD in adipocytes and Peyer’s patch immune cells remains to be understood at a molecular level. An obvious possibility is that, at least, the agonist-dependent element of phosphorylation of Ser^296^/Ser^297^ and Thr^306^/Thr^310^ is mediated by distinct GRKs and that the expression patterns or levels of the GRKs vary between these cell types (45). These and other possibilities will be assessed in future studies. In arrestin-3 interaction studies performed in HEK293 cells alteration of both Thr^306^/Thr^310^ to Ala resulted in only a modest effect on MOMBA-induced recruitment of the arrestin, whereas alteration of both Ser^296^ and Ser^297^ to Ala produced a more substantial effect. However, it would be inappropriate to attempt to correlate this directly with the phosphorylation observed in different native tissues. FFA2 is also expressed in other cell types and tissues, including in pancreatic islets. We have also highlighted previously that the G protein selectivity of agonist-activated FFA2 varies between different cell types and tissues (20). In time, an extensive analysis across a full range of tissues which explores G protein, GRK (and potentially other kinases), and arrestin expression, as well as the integration of functions into cell signalling networks and physiological outcomes, will be required to fully appreciate the implications of tissue-selective FFA2 phosphorylation bar-coding. Despite this, the current studies show clear evidence of how the same ligand-GPCR pairing generates distinct phosphorylation patterns and hence different ‘bar-codes’ in distinct patho-physiologically relevant tissues.

## Experimental

### Materials

Key reagents were obtained from the following suppliers: VECTASHIELD® Antifade Mounting Medium with DAPI (2bscientific, H-1200), anti-HA affinity matrix (Roche, 11815016001), cOmplete™ ULTRA Tablets, Mini, EASYpack Protease Inhibitor Cocktail (Roche, 5892970001), PhosSTOP (Roche, 4906837001), NuPAGE™ MOPS SDS Running Buffer (Thermofisher Scientific, NP0001), NuPAGE™ 4 to 12%, Bis-Tris, 1.0–1.5 mm, Mini Protein Gels (Thermofisher Scientific, NP0321BOX), Tris-Glycine Transfer Buffer (Thermofisher scientific, LC3675), Laemmli buffer (Sigma, 53401-1VL), Nitrocellulose membrane (Thermofisher Scientific, 88018), TSA® tetramethylrhodamine (TMR) detection kit (AKOYA Biosciences, SKU NEL702001KT, Tissue culture reagents (ThermoFisher Scientific). Molecular biology enzymes and the nano-luciferase substrate NanoGlo were from Promega. Polyethylenimine (PEI) (linear poly(vinyl alcohol) [MW 25,000]) (Polysciences). Lambda protein phosphatase (LPPase) was from New England BioLabs. 4-Methoxy-3- methyl benzoic acid (MOMBA) was from Flourochem (018789), whilst (*S*)-3-(2-(3- chlorophenyl)acetamido)-4-(4-(trifluoromethyl)phenyl)butanoic acid (CATPB) and compound 101was from Tocris Bioscience.

### Mutagenesis of DREADD-FFA2-eYFP construct

Site specific mutagenesis on the hFFA2-DREADD-eYFP construct was performed using the Stratagene QuikChange method (Stratagene, Agilent Technologies) and as described in Marsango *et al.,* (31). Primers utilised for mutagenesis were purchased from Thermo Fisher Scientific.

### Cells maintenance, transfection and generation of cell lines

Cell culture, transfection and generation of stable cell lines were carried on as described previously (30).

### Cell lysate preparation

Cell lysates were generated from either the various Flp-In™ T-REx 293™ cells following 100 ng/ml doxycycline treatment or from HEK293T cells following transient transfection, to express C-terminally eYFP-tagged hFFA2-DREADD receptor constructs (hFFA2-DREADD- eYFP) and prepared as described previously (31).

### Cell lysate treatment

To remove phosphate groups, immunocomplexes were treated with LPPase at a final concentration of 10 unit/μl for 90 min at 30 °C before elution with 2 × SDS-PAGE sample buffer.

### Receptor immunoprecipitation and immunoblotting assays

eYFP-linked receptor constructs were immunoprecipitated from prepared cell lysates using a GFP-Trap kit (Chromotek) and immunoblotted as described previously (31). Membranes were incubated with primary antibodies overnight at 4 °C. After overnight incubation the membrane was washed and incubated with secondary antibodies for 2h at room temperature.

### Immunocytochemistry

Flp-In T-REx 293 cells expressing hFFA2-DREADD-eYFP or hFFA2-eYFP were seeded on poly-D-lysine-coated 13 mm round coverslips in 24-well plates and performed as described previously (44).

### Bioluminescence resonance energy transfer-based arrestin-3 recruitment assay

BRET-based arrestin-3 recruitment assay was performed as described in (31). HEK293T cells were transiently transfected with hFFA2-DREADD-eYFP (or each of the indicated DREADD-phosphodeficient (PD) mutants) and arrestin-3 fused to nano-luciferase at a ratio of 1:100. Where indicated, cells were pre-treated with 10 µM compound 101 for 30 mins at 37 °C.

### Tissue dissection

To establish the phosphorylation status of hFFA2-DREADD-HA or hFFA2-HA different tissues from hFFA2-HA, hFFA2-DREADD-HA and the corresponding CRE-MINUS mice were dissected. Following cervical dislocation, the entire colon, Peyer’s patches, mesenteric lymph nodes and adipose tissues were quickly removed and placed in ice-cold Krebs-bicarbonate solution (composition in mmol/L: NaCl 118.4, NaHCO_3_ 24.9, CaCl_2_ 1.9, Mg_2_SO_4_ 1.2, KCl 4.7, KH_2_PO_4_ 1.2, glucose 11.7, pH7.0) supplemented with protease and phosphatase inhibitors. To obtain colonic epithelium preparations, the colon was cut longitudinally and pinned flat on a sylgard-coated Petri dish (serosa down) containing ice-cold sterile oxygenated Krebs solution. The epithelium was gently removed using fine forceps.

### Tissue stimulation

Dissected tissue samples were transferred to warm (30-35°C) Krebs-bicarbonate solution perfused with 95% O_2_, 5% CO_2_ and supplemented with protease and phosphatase inhibitors. Tissue samples were challenged with MOMBA (100µM) (for hFFA2-DREADD-HA) or C3 (2-10mM) (for hFFA2-HA) for 20-30min. For antagonist treatment, tissue samples were incubated with CATPB (10µM) for 30min prior to treatment with MOMBA/C3. Samples intended for western blot analysis were immediately removed and frozen at -80°C. For immunohistochemical analysis tissues were fixed using 4% paraformaldehyde (containing phosphatase inhibitors) for 30min-2h at room temperature.

### Tissue lysate preparation

Frozen samples were homogenised in RIPA buffer (composition in mmol/L: Tris (base) 50mM, NaCl 150mM, sodium deoxycholate 0.5%, Igepal 1%, SDS 0.1%, supplemented with protease phosphatase inhibitor tablets). The samples were further passed through a fine needle (25G) and centrifuged for 15min at 20000g (4°C). Protein quantification was performed using a BCA protein assay kit (ThermoFisher Scientific).

### HA tagged-receptor immunoprecipitation and western blot analysis of tissue samples

HA-tagged receptors were immunoprecipitated overnight at 4°C from the tissue lysates using anti-HA Affinity Matrix beads from rat IgG_1_ (Roche). HA tagged-receptor complexes were centrifuged (2000g for 1min) and washed three times in RIPA buffer. Immune complexes were resuspended in 2x Laemmli sample buffer at 60°C for 5 min. Following centrifugation at 20000g for 2 min, 25 μl of sample was loaded onto an SDS-PAGE on 4- 12% Bis-Tris gel. The proteins were separated and transferred onto a nitrocellulose membrane. Non-specific binding was blocked using 5% bovine serum albumin (BSA) in Tris-buffered saline (TBS, 50 mM Tris-Cl, 150 mM NaCl, pH 7.6) supplemented with 0.1% Tween20 for 1hr at RTP. The membrane was then incubated with appropriate primary antibodies in 5% BSA in TBS- Tween20 overnight at 4°C. Subsequently, the membrane was washed and incubated for 2h with secondary antibodies. After washing (3 × 10 min with TBS-Tween), proteins were visualised using Odyssey imaging system.

### Immunocytochemistry

IHC was performed as previously described by (20). Briefly fixed tissues were embedded in paraffin wax and sliced at 5µM using a microtome. Following deparaffinisation and antigen retrieval, sections were washed in PBS containing 0.3% Triton X-100. Non-specific binding was blocked by incubating sections for 2h at RTP in PBS + 0.1% Triton-X +1%BSA +3% goat serum. Subsequently, sections were incubated in appropriate primary antibodies overnight at 4°C. Sections were washed three times in PBS-Triton X-100 and incubated for 2h at RTP with species specific fluorescent secondary antibodies in the dark. Following three washes, sectioned were mounted with VECTASHIELD Antifade Mounting Medium with DAPI. All images were taken using a Zeiss confocal microscope with Zen software.

### Amplification of HA signal

Signal amplification was performed as previously described by Barki et al., (28). Briefly, sections were incubated with rat anti-HA primary antibody overnight at 4°C, followed by overnight incubation in biotinylated secondary antibody. Immunolabeling was visualised using a TSA tetramethylrhodamine (TMR) detection kit tyramide according to manufacturer’s protocol.

### Antibodies and antisera

The rabbit phospho-site-specific antisera pSer^296^/pSer^297^-hFFA2 and pThr^306^/pThr^310^-hFFA2 were developed in collaboration with 7TM Antibodies GmbH.

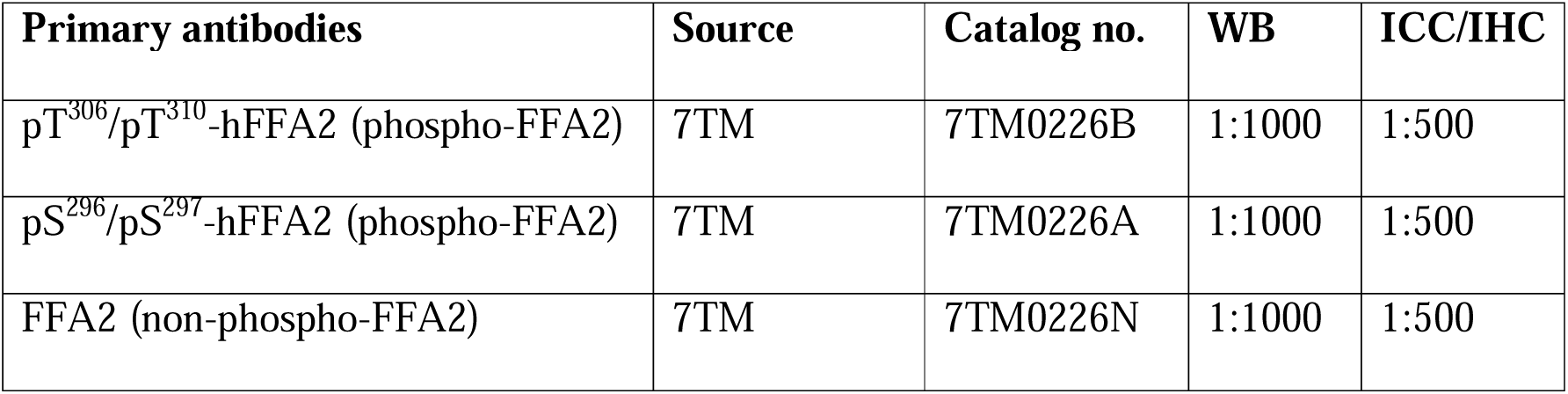
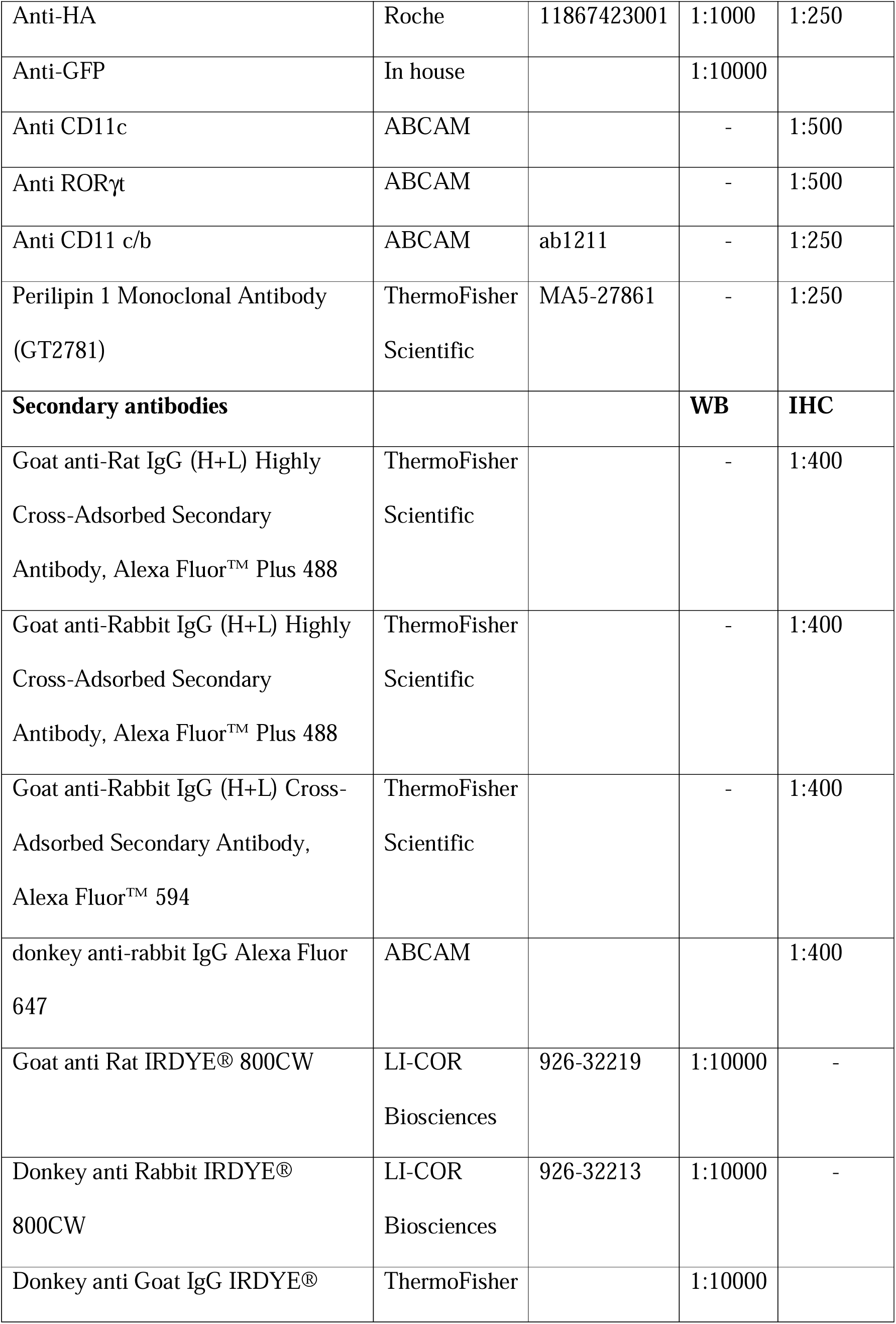
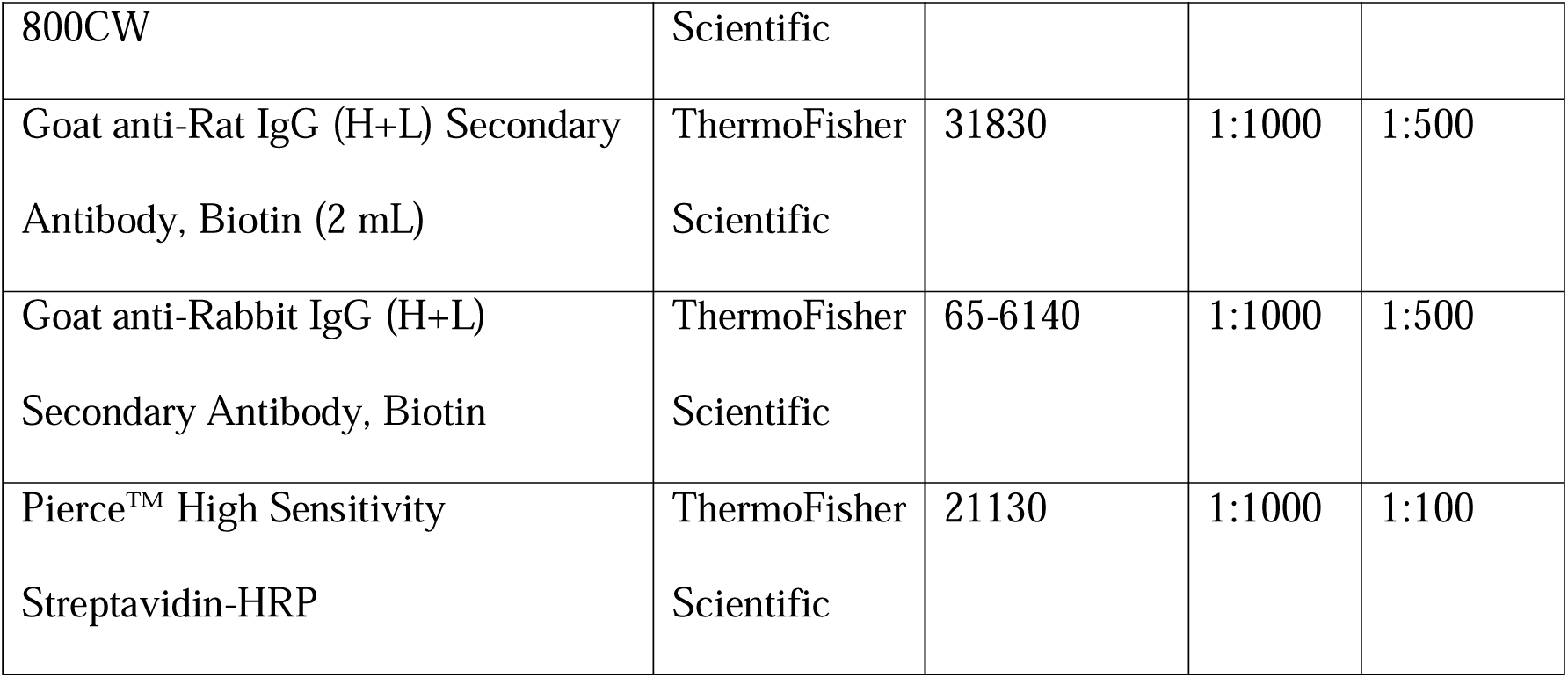

### Treatment, membrane preparation and mass spectrometry analysis of hFFA2- DREADD

Flp-In T-REx 293 cells harboring hFFA2-DREADD-eYFP were cultured to confluence. Receptor expression was induced with 100 ng/mL of doxycycline for 24 h, followed by 5 min stimulation at 37 °C with sorbic acid or MOMBA (100μM). The cells were washed with ice-cold PBS, and TE buffer containing protease and phosphatase inhibitor cocktails was added to the dishes followed by transfer of cells into pre-chilled tubes. The cells were homogenised and centrifuged at 1500 rpm for 5 min at 4 °C. The supernatant was further centrifuged at 50,000 rpm for 45 min at 4 °C. The resulting pellet was solubilised in RIPA buffer [50 mM Tris-HCl, 1 mM EDTA, 1 mM EGTA, 1% (v/v) Triton X-100, and 0.1% (v/v) 2- mercaptoethanol (pH 7.5)] and the protein content was determined using a BCA protein assay kit. hFFA2-DREADD-eYFP was immunoprecipitated using GFP-Trap^®^ Agarose resin (Proteintech) followed by separation on a 4-12% polyacrylamide gel. The gels were stained with Coomassie blue and the bands corresponding to hFFA2-DREADD-eYFP (∼ 75 kDa) were excised. Gel pieces were destained in 50% EtOH and subjected to reduction with 5mM DTT for 30min at 56 °C and alkylation with 10mM iodoacetamide for 30min before digestion with trypsin at 37 °C overnight. The peptides were extracted with 5% formic acid and concentrated to 20 uL. The peptides were subjected to phosphopeptide enrichment using TiO_2_ beads packed into a StageTip column, and phosphorylated peptides were then eluted, dried, and then injected on an Acclaim PepMap 100 C18 trap and an Acclaim PepMap RSLC C18 column (ThermoFisher Scientific), using a nanoLC Ultra 2D plus loading pump and nanoLC as-2 autosampler (Eksigent).

Peptides were loaded onto the trap column for 5 min at a flow rate of 5 ul/min of loading buffer (98% water/2% ACN/ 0.05% trifluoroacetic acid). Trap column was then switched in- line with the analytical column and peptides were eluted with a gradient of increasing ACN, containing 0.1 % formic acid at 300 nl/min. The eluate was sprayed into a TripleTOF 5600+ electrospray tandem mass spectrometer (AB Sciex Pte. Foster City, U.S.A.) and analysed in Information Dependent Acquisition (IDA) mode, performing 120 ms of MS followed by 80 ms MS/MS analyses on the 20 most intense peaks seen by MS.

The MS/MS data file generated was analysed using the Mascot search algorithm (Matrix Science Inc., Boston, MA, U.S.A.), against SwissProt as well as against our in-house database to which we added the protein sequence of interest, using trypsin as the cleavage enzyme. Carbamidomethylation was entered as a fixed modification of cysteine and methionine oxidation, and phosphorylation of serine, threonine and tyrosine as variable modifications. The peptide mass tolerance was set to 20 ppm and the MS/MS mass tolerance to ± 0.1 Da.

Scaffold (version Scaffold_4.8.7, Proteome Software Inc) was used to validate MS/MS-based peptide and protein identifications. Peptide identifications were accepted if they could be established at greater than 20.0% probability. Peptide Probabilities from X! Tandem and Mascot were assigned by the Scaffold Local FDR algorithm. Peptide Probabilities from X!

Tandem were assigned by the Peptide Prophet algorithm with Scaffold delta-mass correction. Protein identifications were accepted if they could be established at greater than 95.0% probability and contained at least two identified peptides.

## Data Availability

The hFFA2-DREADD-HA, hFFA2-HA and corresponding CRE-MINUS mouse lines are available upon request to either ABT or GM. The mass spectrometry proteomics data have been deposited to the ProteomeXchange Consortium via the PRIDE [46] partner repository with the dataset identifier PXD042684. All other data is freely available from G.M. or A.B.T. or or through the University of Glasgow’s online data repository.

## Acknowledgments and Funding Sources

These studies were supported by the Biotechnology and Biosciences Research Council U.K., grants numbers BB/X001814/1 and BB/S000453/1 (to GM and ABT).

## Supplementary Information

**Supplementary Figure 1.**
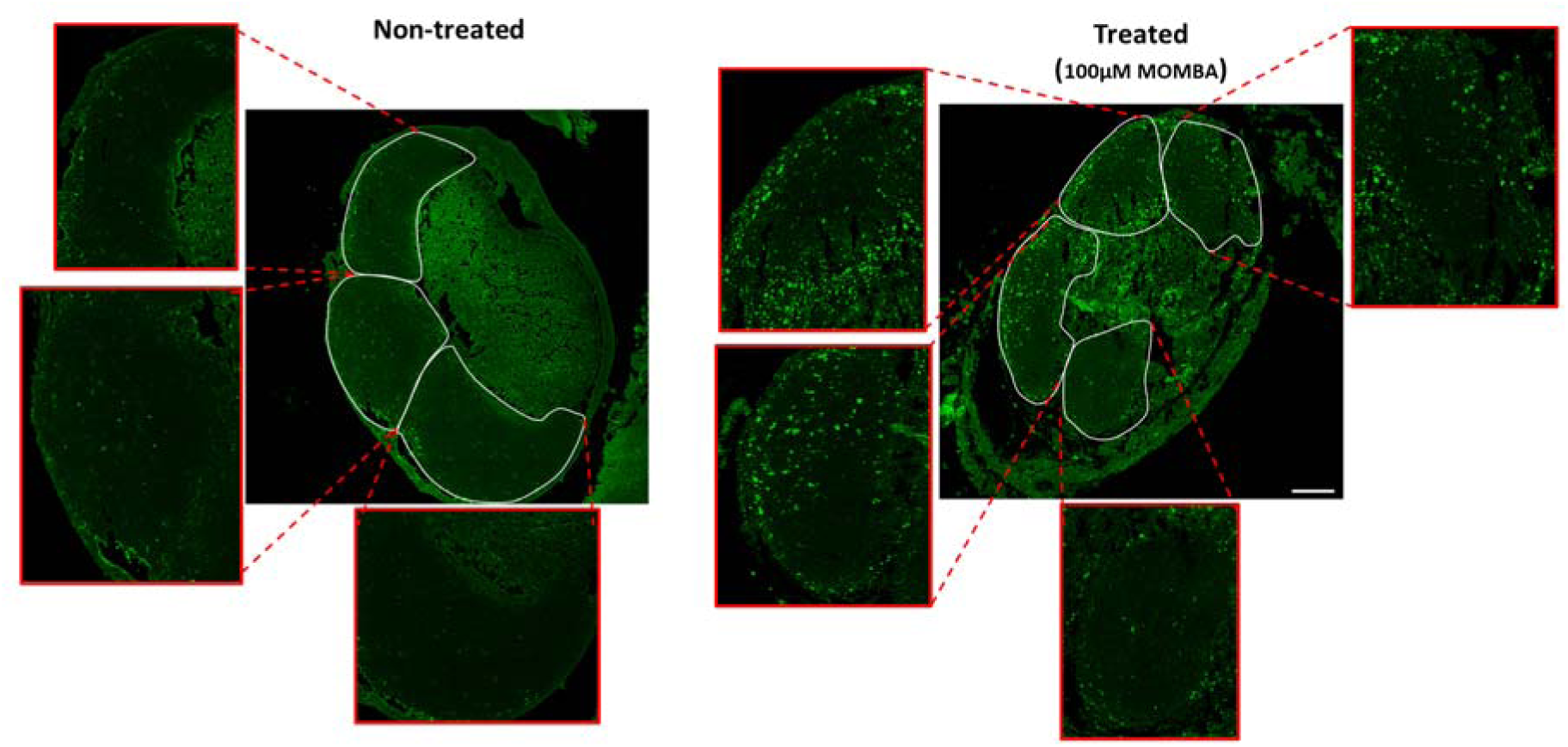
hFFA2-DREADD-HA becomes phosphorylated at Thr ^306^ /Thr ^310^ in immune cells within Peyer’s patches. Experiments akin to those of Figure 5F were conducted using the pThr ^306^ /pThr ^310^ hFFA2 antiserum. The figure illustrates MOMBA-induced phosphorylation of pThr ^306^ /pThr ^310^ in immune cells within Peyer’s patches of hFFA2-DREADD-HA expressing mice. Peyer’s patches were either treated with vehicle (**Left panel**) or MOMBA (100 μM) (**Right panel**). Each lymphoid nodule has been expanded to show detailed phosphorylation inside each nodule.

**Supplementary Figure 2.**
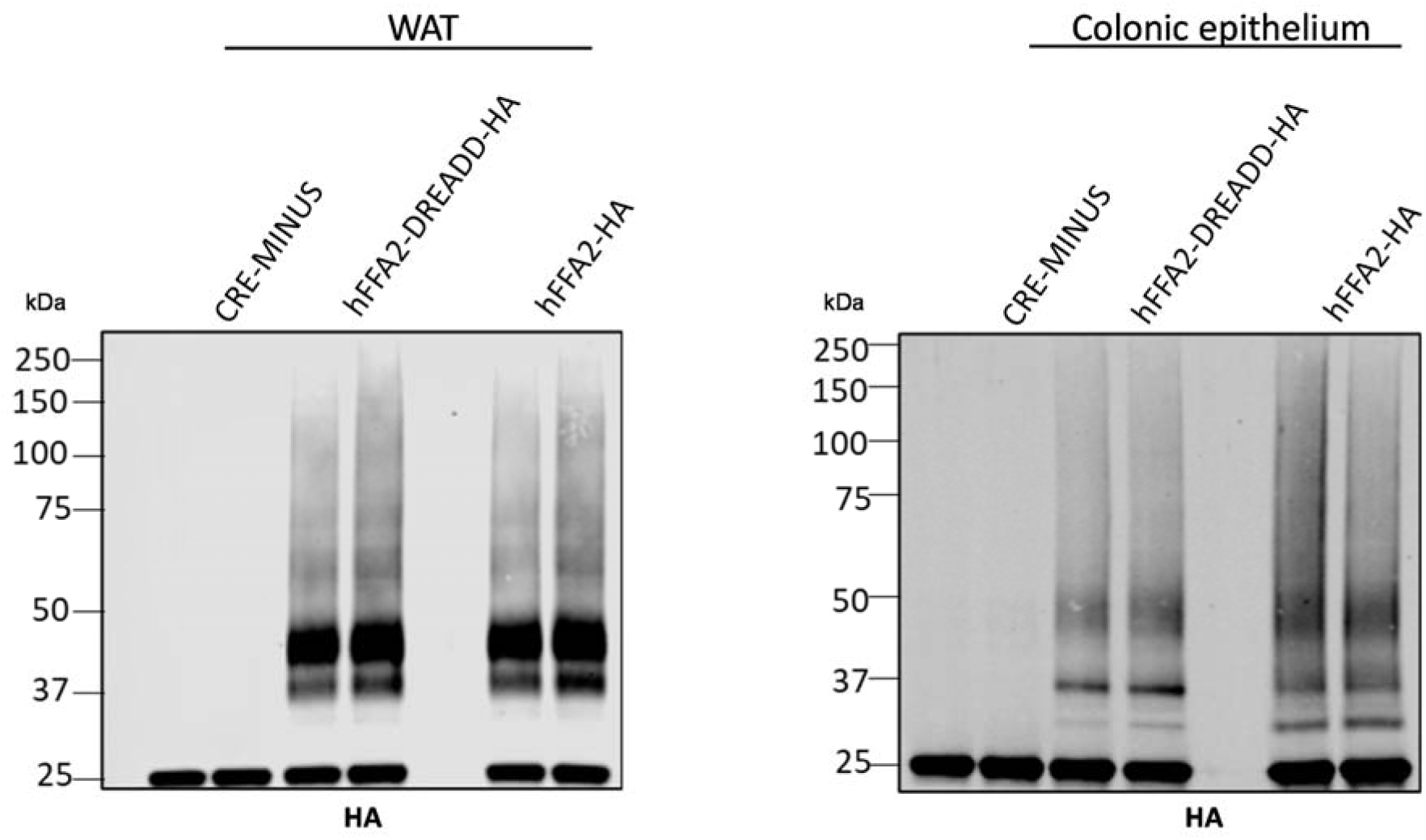
Tissues of transgenic mice express similar levels of hFFA2-HA and hFFA2-DREADD-HA. White adipose (WAT) (**left panel**) and colonic epithelial (**right panel**) tissue was isolated from CRE-MINUS and both hFFA2-DREADD-HA and hFFA2-HA expressing transgenic mice. Anti-HA immunoprecipitations were then performed, and samples resolved by SDSPAGE followed by immunoblotting with anti-HA. For both tissues similar levels and patterns of the receptor proteins were detected from the hFFA2-DREADD-HA and hFFA2-HA expressing mice, whilst these were absent in tissue from CRE-MINUS animals. A representative experiment of 3 is shown.

**Supplementary Figure 3.**
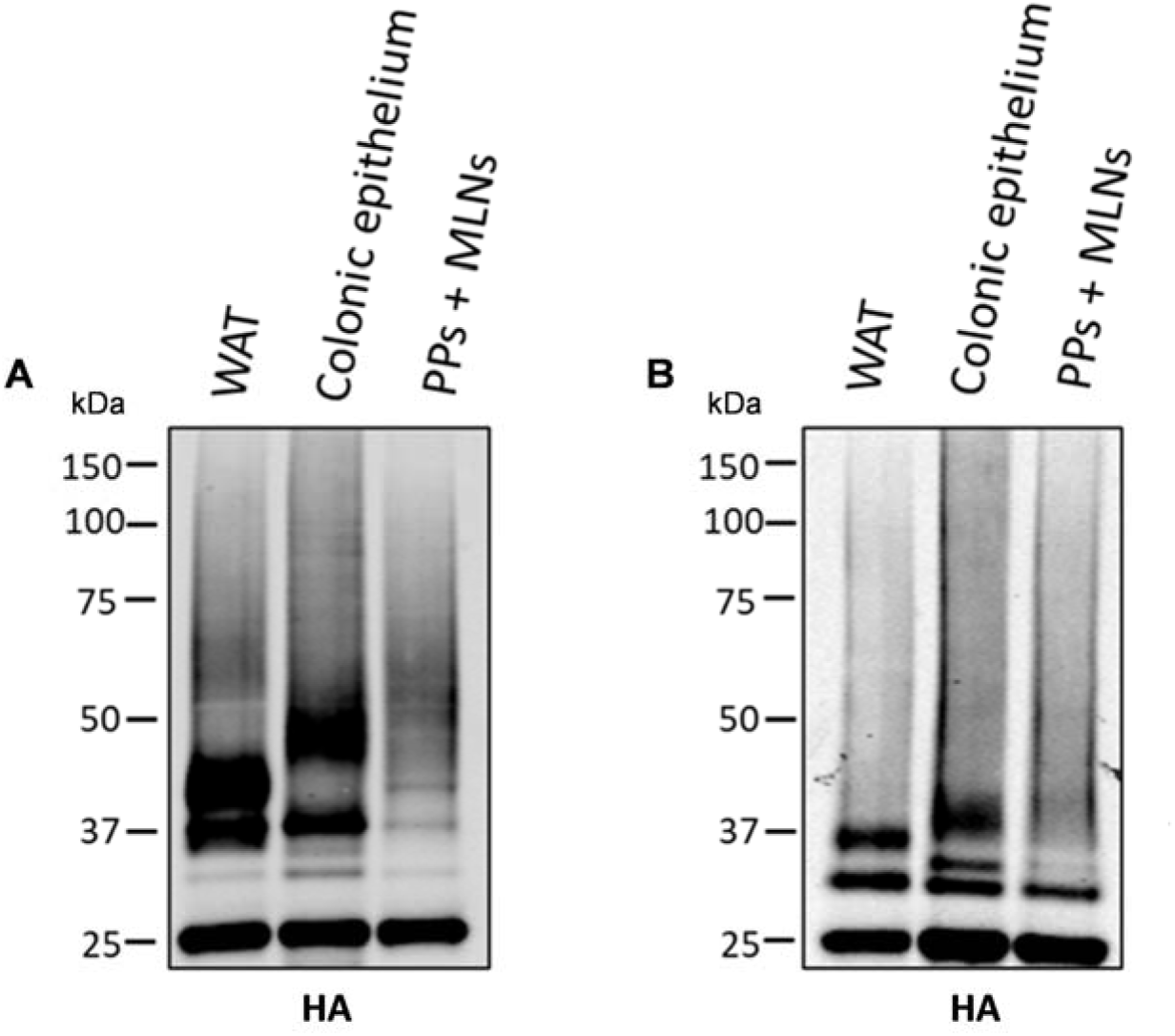
hFFA2-DREADD-HA is present as multiple differentially N-glycosylated species in adipose tissue, immune, and colonic epithelial cells. HA-immunoprecipitated samples from white adipose tissue (WAT), colonic epithelium and Peyer’s patches and mesenteric lymph nodes (PP + MLNs) of hFFA2-DREADD-HA expressing mice were untreated (**A**) or treated with N-glycosidase F to remove N-linked carbohydrate (**B**). They were then resolved by SDS-PAGE and immunoblotted with anti-HA. A representative experiment of 3 is shown.

